# Structural basis for plasmid restriction by SMC JET nuclease

**DOI:** 10.1101/2023.11.03.565488

**Authors:** Florian Roisné-Hamelin, Hon Wing Liu, Michael Taschner, Yan Li, Stephan Gruber

## Abstract

DNA loop-extruding SMC complexes play crucial roles in chromosome folding and DNA immunity. Prokaryotic SMC Wadjet (JET) complexes limit the spread of circular plasmids through DNA cleavage; yet the mechanisms for target recognition are unresolved. We show that artificial DNA circularization renders linear DNA susceptible to JET cleavage. Unlike free DNA, JET cleaves immobilized plasmid DNA at a specific site, the plasmid-anchoring point, showing that the anchor hinders DNA extrusion but not DNA cleavage implying that residual unextruded DNA is cleaved. Structures of plasmid-bound JetABC reveal two presumably stalled SMC motor units that are drastically rearranged from the resting state, together entrapping a U-shaped DNA segment, which is further converted to kinked V-shaped cleavage substrate by JetD nuclease binding. Our findings uncover mechanical bending of residual unextruded DNA as principle for non-self DNA recognition and molecular signature for plasmid cleavage. We elucidate key elements of SMC loop extrusion including motor directionality and the structure of a DNA-holding state.

## Introduction

SMC complexes represent ATP-powered DNA motors that fold DNA segments of megabase size through loop extrusion. They play crucial roles in various cellular processes, including chromosome segregation during cell division (condensin and cohesin in eukaryotes, Smc-ScpAB, MukBEF, and MksBEF in prokaryotes), the regulation of gene expression, DNA repair, and recombination (cohesin and Smc5/6 in eukaryotes) ^1–3^. More recently, emerging evidence suggests that SMC-based complexes are also involved in cellular defence against invasive or selfish genetic elements ^4^. In prokaryotes, the JET complex (short for Wadjet, JetABCD) is a sequence-independent anti-plasmid nuclease ^5–10^, while the human Smc5/6 complex functions as a viral restriction factor ^4,11^. Nevertheless, the role of loop extrusion in DNA immune sensing remains unclear ^8,9^. The process of DNA loop extrusion by the eukaryotic SMC complexes has been observed directly by single-molecule experiments *in vitro* ^12–15^. For bacterial Smc-ScpAB complexes, loop extrusion has been inferred from and characterized through chromosome conformation capture-type experiments *in vivo* ^16^. Yet, the molecular mechanism and structural basis of SMC loop extrusion remain enigmatic and subject to debate, resulting in a range of proposed models ^1,17,18^. The determination of structures of SMC motors bound to physiological DNA substrates represents a crucial goal but also a significant challenge.

The multi-subunit SMC complexes are comprised of an SMC protein dimer (JetC in JET), a kleisin subunit (JetA), and accessory KITE (JetB) or HAWK proteins. SMC proteins feature a long coiled coil “arm” that harbors a “hinge” dimerization domain at one end and an ATP binding cassette “head” domain at the other. Connecting the two SMC proteins, the kleisin subunit forms the distinctive elongated tripartite ring structure that enables stable chromosome binding through DNA entrapment. Upon ATP binding/sandwiching, the head domains engage with each other, disrupting the alignment of SMC arms and facilitating DNA clamping. This process likely involves the capture of a looped DNA segment between the SMC arms by exposing a head-DNA binding surface. The kleisin-associated KITE or HAWK subunits contribute to DNA clamping/segment capture by associating with both DNA and the SMC proteins ^19–23^. ATP hydrolysis is thought to reset the complex into a “DNA holding” configuration where the heads are juxtaposed and the arms are closed ^19^. This pentameric assembly likely constitutes the minimal unit for DNA translocation, here referred to as the “motor unit” ^1–3,17^. Certain SMC complexes form stable dimers (i.e. dimers of the pentameric motor unit) ^24,25,22^. In case of the JET complex (Figure 1A), dimerization is facilitated by homotypic interactions between amino-terminal kleisin JetA sequences ^8,9^. The JetD subunit is exclusive to the JET family of SMC complexes ^6,26^. JetD encompasses rigidly connected amino-terminal arm and CAP domains (denoted as “aCAP”), as well as a flexibly connected carboxy-terminal Toprim domain. This configuration bears similarities to a subunit of an archaeal type II DNA topoisomerase (topoisomerase VI) and the meiotic recombination initiator Spo11 (Figure 1A) ^5,6,8–10,26,27^. JetD serves the nucleolytic function of the JET complex ^8–10^. However, JetD remains inactive in isolation; likely DNA restriction necessitates the alleviation of JetD autoinhibition by JetABC ^8,9^.

**Figure 1:**
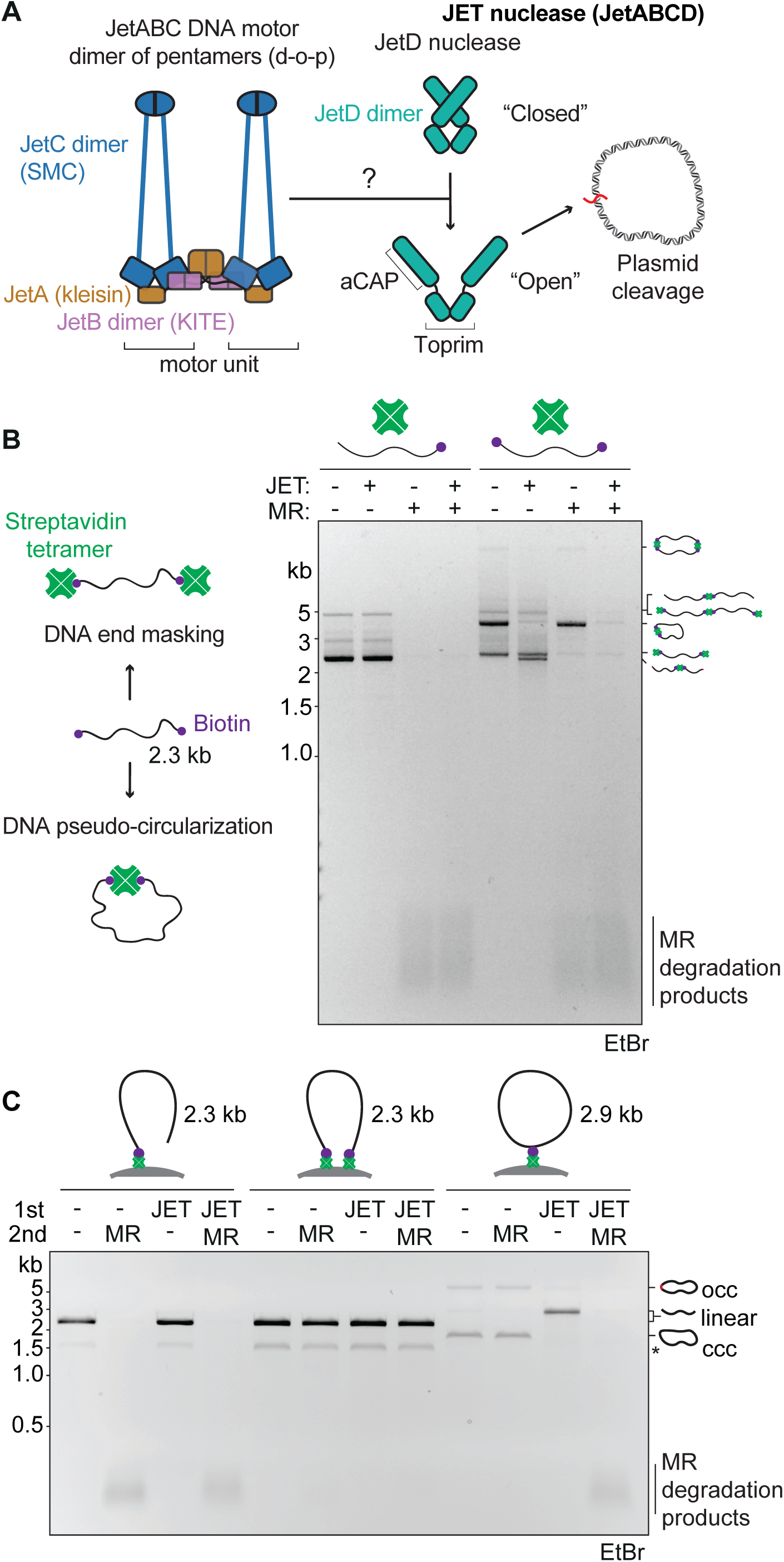
JET cleavage of pseudo-circular DNA. (A) Schematic representation of *E. coli* GF4-3 (Type I) JET nuclease dimer-of-pentamers (d-o-p) in the resting state ^4,8^. (B) Left: Schematic depicting DNA end masking and pseudo-circularization when streptavidin tetramer is added to biotinylated linear DNA. Right: Cleavage assays with JET nuclease (JetABC: 12.5 nM d-o-p, JetD: 25 nM dimer) and MR nuclease (125 nM, tetramer) on DNA species obtained by streptavidin (100 nM monomer) incubation with single or double biotinylated 2.3 kb DNA substrates (7 nM). The resulting products were resolved on a 1% agarose gel containing ethidium bromide (EtBr). (C) Cleavage assay of biotinylated linear DNA anchored onto streptavidin-coated dynabeads. See Figure S2B and Methods for experimental pipeline. Biotinylated DNA circles (ccc, closed covalent circles and occ, open covalent circles) were used to demonstrate JET activity (see also Figure S2). The asterisk indicates a low abundance DNA contaminant obtained by PCR. See also Figures S1 and S2.

JET systems exhibit a specific capacity to restrict smaller (<80-100 kb) circular DNA molecules through DNA cleavage, independently of DNA sequence and helical topology (Figure 1A) ^8,9^. This requires ATP, functional JetC ATPase, and JetD nuclease, giving rise to linear products with some variability in the nature of the DNA ends^6,8,9^. The precise mechanism underlying plasmid restriction by the JET complex remains elusive. It is plausible that the JetABC motors employ a DNA extrusion reaction to discern DNA topology and size ^8,9^. This raises the intriguing question of how the activation of the JetD executor might be coupled to any JetABC DNA extrusion activity ^8,9^. Furthermore, the way the JET DNA motor (or any SMC DNA motor) might navigate DNA-bound proteins, referred to as “roadblocks”, is unclear. Single-molecule experiments have suggested that SMC complexes possess the capability to bypass substantial roadblocks ^13,28^, challenging the notion that DNA entrapment maintain SMC complexes stably associated with DNA and translocating in a directional manner.

Here, we explore the molecular and structural mechanisms governing DNA recognition and cleavage by a type I *Escherichia coli* JET nuclease complex ^6,8^. We show that circular DNA substrates with a single large roadblock prompt the JET nuclease to cleave in proximity to the anchoring point. Conversely, substrates with two large roadblocks impede cleavage, collectively indicating that JET is unable to bypass larger roadblocks and underscoring the necessity for extruding most of the plasmid DNA, albeit not all, prior to initiating cleavage. Through cryo-EM analyses, we unveil the structures of both JetABC and nuclease-defective JetABCD bound to plasmid DNA. These structures reveal configurations of the JET nuclease that are competent for cleavage, featuring an alternative SMC dimer arrangement and a tightly bound DNA U turn. These characteristics appear to arise from DNA motor stalling on the diminishing section of the plasmid DNA. A JetD dimer binds to JetABC-bent DNA and introduces a kink prior to DNA cleavage. In summary, our findings strongly support a model in which JET motors actively survey DNA molecules through loop extrusion to selectively target and restrict smaller circular plasmids.

## Results

### JET cleavage of linear DNA upon artificial circularization

Linear DNA molecules exhibit resistance to cleavage by JET ^8,9^. We wondered whether the JET nuclease detects the absence of DNA ends in circular DNA or senses the continuity of its DNA double helix. We asked whether linear DNA becomes susceptible to cleavage when its DNA ends are masked or when they are artificially linked together to form pseudo-circular DNA (Figure 1B). We employed the binding of streptavidin tetramers to biotin-labeled DNA molecules, which were generated by PCR using 5’ biotinylated oligonucleotide primers. A related approach has recently shown that the endonuclease Rad50/Mre11 (MR) (Figure S1A) recognizes DNA ends even in instances where they are masked, but not when they are linked together to create pseudo-circular DNA ^29^. Incubation of double biotin-labelled DNA molecules with streptavidin generated a mixture of DNA species (Figures 1B and S1B). Most species originating from double-biotin DNA exhibited resistance to RecBCD due to double end masking (Figure S1B). However, only a singular species exhibited resistance to the MR nuclease, which corresponded to monomeric pseudo-circular DNA (Figure 1B) ^29^. Remarkably, while JET nuclease was incapable of cleaving linear DNA even when both ends were concealed, it efficiently cleaved the pseudo-circular DNA (Figure 1B) at largely random positions (Figure S1C). This unequivocally demonstrates that the JET nuclease senses the circular nature of DNA rather than the mere absence of free DNA ends. Species that evaded MR were targets of JET action, and conversely, species that resisted JET were sensitive to MR. Accordingly, dual treatment with MR and JET led to the transformation of all DNA species into short DNA fragments (Figure 1B). These outcomes strongly suggest that the JET nuclease surveys the target DNA to ascertain DNA (pseudo-) circularity prior to triggering DNA cleavage. Conversely, connecting the two DNA ends at a larger distance (via binding to a large polystyrene bead) did not render DNA susceptible to JET cleavage, despite becoming protected from MR-induced degradation (Figure 1C). This implies that a continuous DNA translocation track is required for recognition by the JET nuclease.

### JET cleavage at the DNA anchoring sites of large roadblocks

The above results indicate that JET activity is robust even when the double-stranded DNA contains structural perturbations (in the form of streptavidin-biotin junctions), as long as DNA retains its (pseudo-) circular nature. Similar outcomes were observed for circular DNA substrates harboring interruptions in the form of a 10 nt single-stranded DNA flap or a 68 nt single-stranded DNA gap (Figure S1D-E). We wondered whether larger obstacles on DNA would hinder DNA cleavage by representing impassable barriers for plasmid DNA extrusion. Our initial approach involved utilizing streptavidin-biotin linkage to tether plasmid DNA to large spherical polystyrene beads (dynabeads; 2.8 µm in diameter). We observed minimal effects on DNA cleavage (and mild effects on cleavage site distribution) (Figures 1C and S2A-D), potentially indicating that JET was able to bypass even substantial roadblocks as recently suggested by single-molecule imaging work on other SMC complexes (that also employed non-covalent DNA-bead attachments) ^28^. However, it is conceivable that the slight effects observed here, as well as in the single-molecule work, arose from transient leakage of DNA from the beads (Figure S2E). Indeed, we observed low but noticeable release of biotin-DNA from the beads during the experiments, which was strongly influenced by the biotin moiety’s position and DNA’s shape (Figure S3A-D).

To firmly rule out this possibility, we established a more rigorous approach that exploits SNAP-tag technology to establish an uninterrupted covalent link between dynabeads and DNA (modified with SNAP-ligand benzylguanine, BG using a gap-filling approach ^30^) (Figures S2A and S4A-C, see Methods). We found that a large fraction of DNA circles was cleaved by JET nuclease, even when covalently linked to dynabeads (Figure 2A). This observation confirms that the presence of a large roadblock does not necessarily halt DNA cleavage by JET. Notably, subsequent treatment with the single-cutting restriction enzyme ScaI resulted in distinct bands instead of a smear, indicating JET cleavage occurred at specific positions (Figure 2B). The position of DNA cleavage corresponded with the position of the SNAP-bead anchor indicating that it occurred at or in very close proximity to the roadblock anchor. Unlike the outcomes from the biotin-streptavidin experiments (Figure S2D), no DNA smearing was observed in this case, strongly suggesting that all cleavage by JET happened near the anchor point (Figure 2B). In contrast, when utilizing a smaller roadblock, SNAP-tag not chemically tethered to dynabeads, a pattern of random DNA cleavage was observed (Figure S4D). This contrast implies that the JET nuclease can efficiently navigate through smaller (∼10 nm range) DNA roadblocks that remain continuously attached to DNA but not when the roadblocks are significantly larger (within the micrometer range). If large roadblocks indeed block DNA extrusion but not DNA cleavage, then complete extrusion cannot be a prerequisite for DNA cleavage, suggesting that cleavage occurs on a (residual) unextruded segment of the plasmid (Figure 2D).

**Figure 2:**
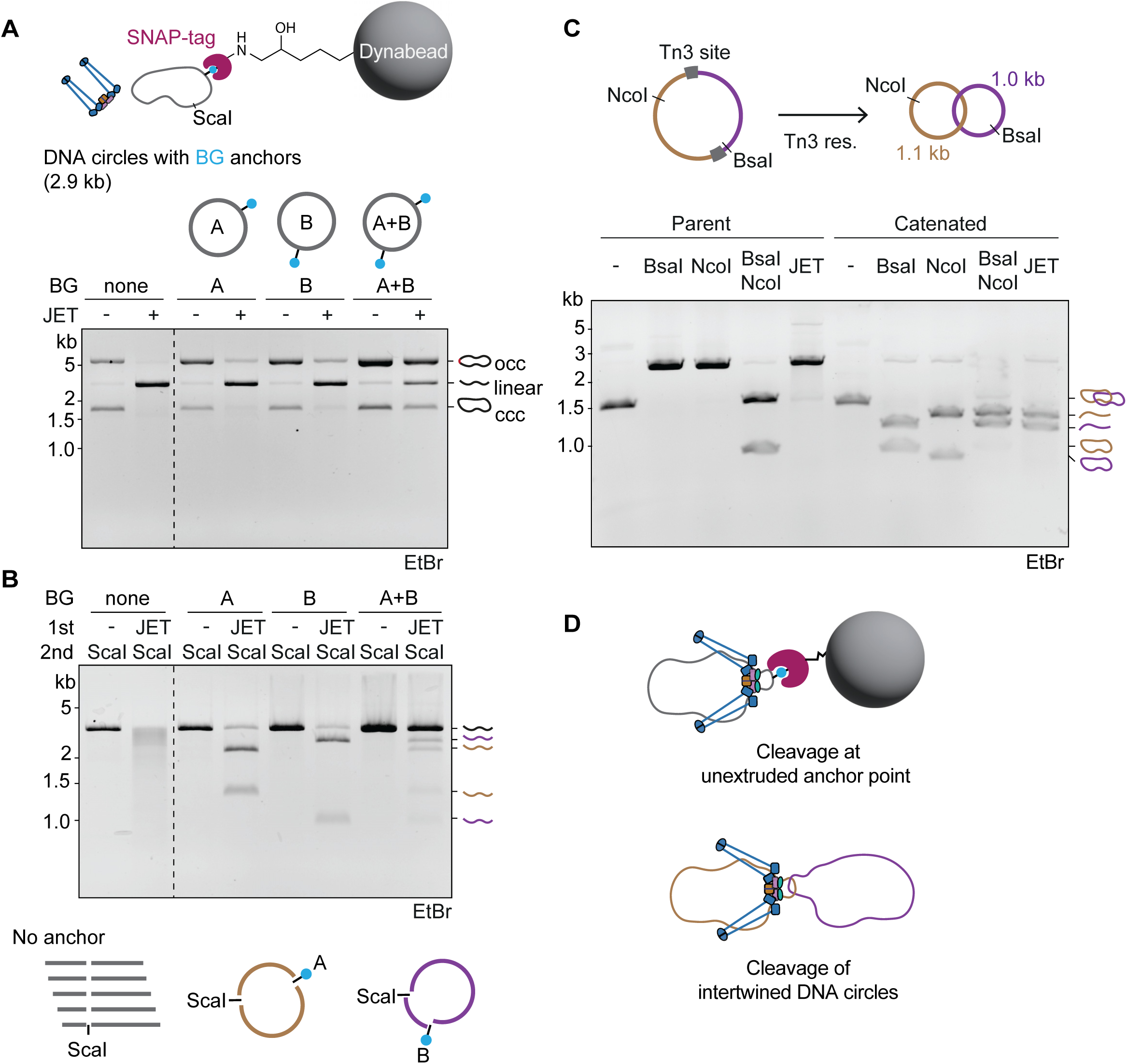
JET cleavage of circular DNA with roadblocks. (A) Top: Schematic of DNA covalently anchored onto dynabeads. See Figure S2A for a plasmid map containing the modification sites, and Figure S4C for reaction pipeline. Bottom: JET cleavage of bead- anchored DNA. We note that a population of BG-DNA circles migrated slower than closed covalent circles (ccc) — likely, these open covalent circles (occ) were generated by DNA strand breakage during prolonged treatment for DNA anchoring. We adopted a gap-filling approach ^30^ to generate covalently closed 2.9 kb DNA circles containing one or two SNAP ligands, benzylguanine groups (BG), attached to selected thymidine nucleobases (see also Figures S2A, S4A). The labelled DNA was then reacted with purified SNAP-tag protein (Figure S4B), and further anchored to dynabeads through chemical cross- linking via amino-reactive epoxy chemistry. Uncoupled DNA was removed by washing before treatment with JET nuclease. For gel analysis, the DNA covalently coupled to the beads was eventually released by digesting the SNAP-tag protein using proteinase K (Figure S4C). (B) JET cleavage site distribution. Top: Agarose gel depicting fragmentation obtained from ScaI post- treatment of JET cleaved products. Bottom: Schematics explaining the outcomes above. (C) Top: Schematic of a DNA catenane generated by Tn3 site-specific recombination. Bottom: Agarose gel showing DNA cleavage activity of JET (12.5 nM d-o-p) on catenated DNA circles (8.1 nM). (D) Schematics depicting a putative JET cleavage state when encountering a large roadblock (top) or intertwined DNA (bottom). See also Figures S2, S3 and S4.

Consistent with this notion, we observed a marked reduction in the cleavage of plasmids harboring two SNAP ligands for bead anchoring, even when subjected to extended periods of incubation (Figures 2A and S4E). We note that a pool of DNA remained susceptible to cleavage even at earlier reaction times, possibly emerging from DNA with only a single covalent link to dynabeads owing to incomplete (SNAP or epoxy) coupling (Figures 2A and S4E). A likely interpretation is that a double roadblock prevents JetABCD motor units from converging on the plasmid DNA, thus impeding DNA cleavage. In the same vein, we found that catenated DNA circles, generated through site-specific recombination from a precursor plasmid ^31^, were efficiently cleaved by JET nuclease (Figure 2C). In this case, the complete extrusion of a given DNA circle is expected to be inhibited by the presence of the interlinking DNA molecule. We propose that the interlinked DNA (just like the anchor for a single bead roadblock) is accommodated together with the unextruded DNA in a cleavage-proficient JET structure (Figure 2D).

### Cryo-EM structure of plasmid-bound JetABC

To visualize how the DNA motor units may converge on plasmid DNA, we performed cryogenic electron microscopy (cryo-EM). Initially, we excluded the nuclease subunit JetD and incubated JetABC with plasmid DNA (pDonor, 1.8 kb) for 10 minutes at room temperature in the presence of ATP prior to grid freezing. This is expected to provide sufficient time for DNA sensing by JET and its priming for cleavage (given that JetABC concentration, rather than JetD, is rate-limiting for cleavage ^8^). This approach yielded a structure of the dimeric core of plasmid-borne JetABC, with an overall resolution of 4.8 Å (Figures 3A and S5; Table S1, Methods). As with the previously reported “resting state” structure (lacking DNA) ^8,9^, certain segments, namely the more distal regions of the JetC arms and the hinge, were not resolved in our map, likely due to inherent flexibility (attempts at local refinement were not successful, see Methods). The resulting map revealed a novel JetABC dimer geometry that is strikingly distinct from the resting state ^8,9^, as well as from the MukBEF complex ^22^. This altered geometry originates from a distinct configuration of the JetA kleisin dimer, which we discuss further below using a higher-resolution structure and a corresponding model (Figures 4-5 and S6-8). Remarkably, we find that both DNA motor units engage with a tightly bent U-shaped DNA molecule with a ∼60 bp DNA half circle at its center. This looping is apparently not resulting from DNA binding to a curved surface, but rather emerges from external mechanical constraints imposed upon the DNA double helix by the adjacent JetABC dimer (Figure 3A). How the constraints are generated is not immediately clear, but DNA pulling by JET DNA motor activity seems like the most plausible scenario.

**Figure 3.**
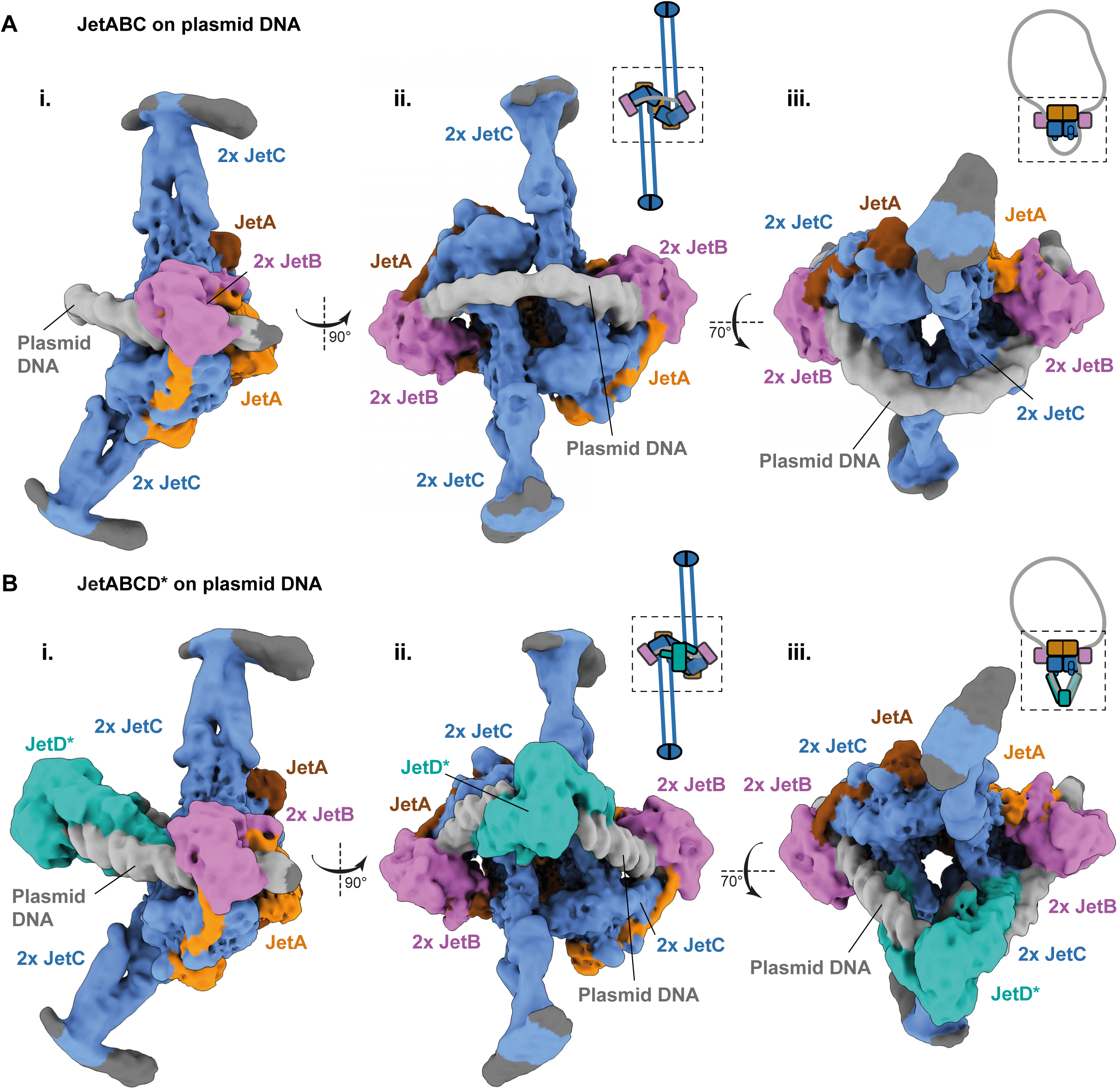
Cryo-EM structures of plasmid-borne JetABC and JetABCD*. (A) Cryo-EM structure of plasmid-borne JetABC (map locally filtered in cryoSPARC, overall resolution: 4.8 Å in three orientations (i, ii, iii). JetA, JetB, JetC and DNA are shown in yellow, purple, blue, and grey colors, respectively. (B) Cryo-EM structure of plasmid-borne JetABCD* (map locally filtered in cryoSPARC, overall resolution: 4.35 Å) shown in the same orientations as in (A) (i, ii, iii). JetD* corresponds to the JetD(E248A) mutant. Coloring as in (A) with JetD shown in turquoise colors. See also Figures S5, S6 and Table S1.

**Figure 4.**
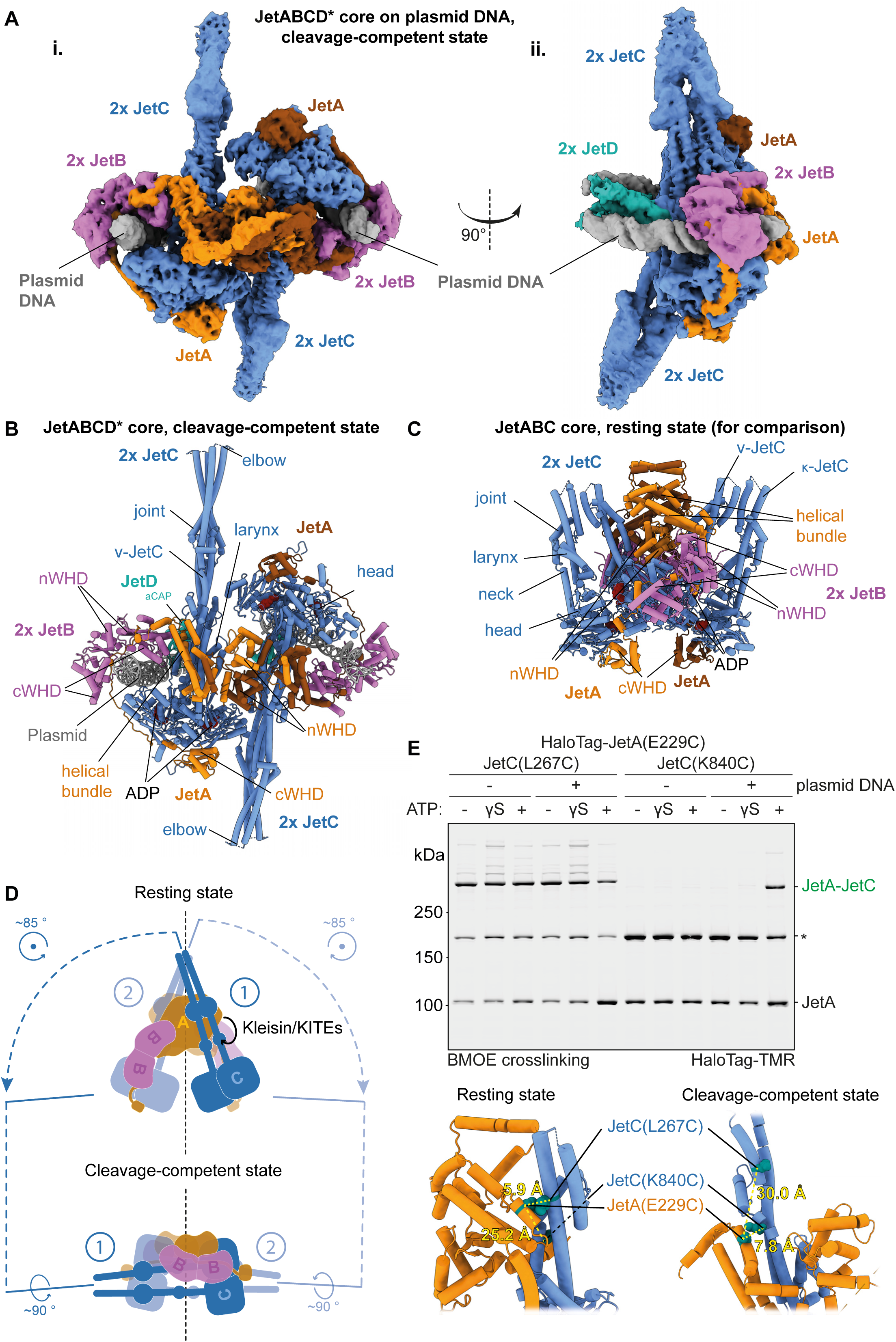
Architecture of the JetABCD* core in the cleavage-competent state. (A) Cryo-EM structure of plasmid-borne JetABCD* core (including JetC heads and head-proximal coiled coils, JetB dimers, a JetA dimer, and JetD aCAP; overall resolution: 4.2 Å). Based on data shown in Figure 3B (using non-uniform refinement with a mask on the core). Two orientations are shown (i, ii). (B) Model of the JetABCD* core (see Methods; Figure S7; Table S1); oriented as in (A i). (C) Model of the JetABC core in the resting state (PDB:8BFN) ^8^ shown for comparison. (D) Schematic illustration of motor dimer geometry in the resting (top panel) and the cleavage- competent state (bottom panel). The motors are aligned via the JetA nWHD dimer located at the dyad axis (shown as dashed vertical lines). Transformations of the JetC dimers and the JetA helical bundle/JetB dimer are indicated by blue and black arrows, respectively. (E) Cysteine cross-linking of JetA/JetC interface residues. Top: Imaging of TMR-labelled JetA-HaloTag after BMOE cross-linking of JetABC in the indicated conditions. A-C: JetA-JetC cross-linked product; A: JetA. Of note, we also readily detected JetA-JetA cross-linked product (indicated by asterisk), stemming from JetA(E229C) cross-linking, possibly between dimers of pentamers. Alternatively, the cross-linking may result from an unknown (intermediary) conformation(s) of the complex. Bottom: Position of the engineered cysteine residues shown as balls in green colors in the models for the resting (PDB: 8BFN) and cleavage-competent states. The yellow dashed lines indicate the Cα-Cα distances between selected cysteine pairs. See also Figures S6, S7, S8, S9, S10 and Table S1.

**Figure 5.**
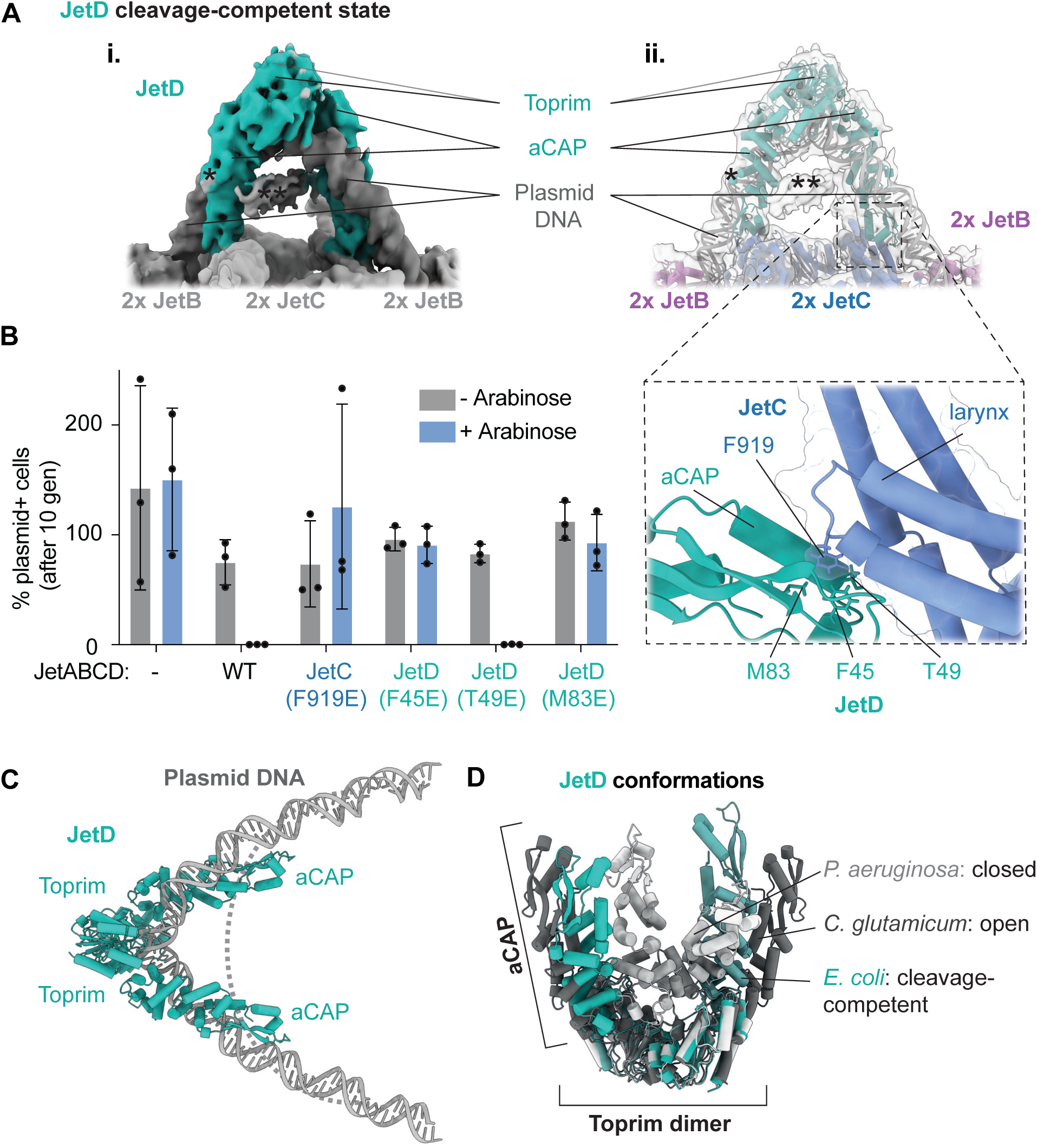
JetD architecture in the cleavage-competent state. (A) Local map of a JetD dimer bound to plasmid DNA (i) (see Table S1; Methods; Figures S6 and S11) and AlphaFold2-predicted models of JetD domains and B-form DNA fitted into the map (ii). The interface between JetD and JetC is marked by the box and shown in larger magnification in (B). Of note, extra density not unambiguously assigned was marked by asterisks. The extra density labelled by a single asterisk may correspond to a short JetB amino-terminal helix that was predicted to bind to JetD aCAP ^9^, while the other density may originate from remaining flexible JetB sequences. (B) Left: Plasmid restriction by mutant JET. Graph showing the percentage of plasmid (pBAD)- containing *E. coli* cells in a cell population after ten generations without selection with or without JET induction by arabinose addition. Means and standard deviations from three independent experiments are shown. Right panel: Model of the JetD(aCAP)-JetB(larynx) interface depicting residues tested by mutagenesis are stick representation (JetABCD* core model, Figure 4B), see also Figure S9. The JetC protein surface is displayed in semi-transparent blue colors. (C) A model of the JetD dimer bound to plasmid DNA (as described in panel A) shows V-shaped DNA with the apex located at the Toprim domains. The dashed curve represents the DNA curvature in the absence of JetD. Other parts of the structure are omitted for clarity. (D) Comparison of JetD conformations. The *E. coli* JetD dimer in turquoise colors as described in (A) was aligned via the Toprim domain dimer with the model of *Pseudomonas aeruginosa* JetD in a closed conformation (PDB:7TIL) ^9^ in light grey colors and the model of *Corynebacterium glutamicum* JetD in an open conformation (PDB:8B7F) ^10^ in dark grey colors. See also Figures S6, S9, S11 and Table S1.

Each motor unit entraps the plasmid DNA molecule in the JetA kleisin compartment. The JetB KITE dimer engages with DNA in a manner analogous to the DNA clamping state seen in MukBEF ^22^. However, the DNA does not traverse between the SMC arms as is the case in the DNA clamping/DNA segment capture state, revealing a “DNA holding” SMC configuration as previously inferred from cysteine cross-linking experiments ^19,23,32,33^ (Figure 3A, see also Figure 6B). We note that in a structural model (described further below; Figures 4 and S6-7) several positively charged residues of JetB are close to the DNA phosphodiester backbone (Figure S9A), mutation of which abolished plasmid restriction activity *in vivo* (Figure S9C), suggesting that they are important for DNA sensing or processing. The structure represents a “post-hydrolysis” configuration as previously reported for the resting state ^8^ (Figure 3A, see also Figure S8C). In this configuration, the ADP-bound heads are juxtaposed yet not engaged, while the arms are closed. This configuration implies that the SMC compartment is devoid of DNA, altogether providing strong evidence for genuine topological DNA entrapment. Stable DNA entrapment offers a plausible explanation for the observed difficulty in bypassing large roadblocks.

**Figure 6.**
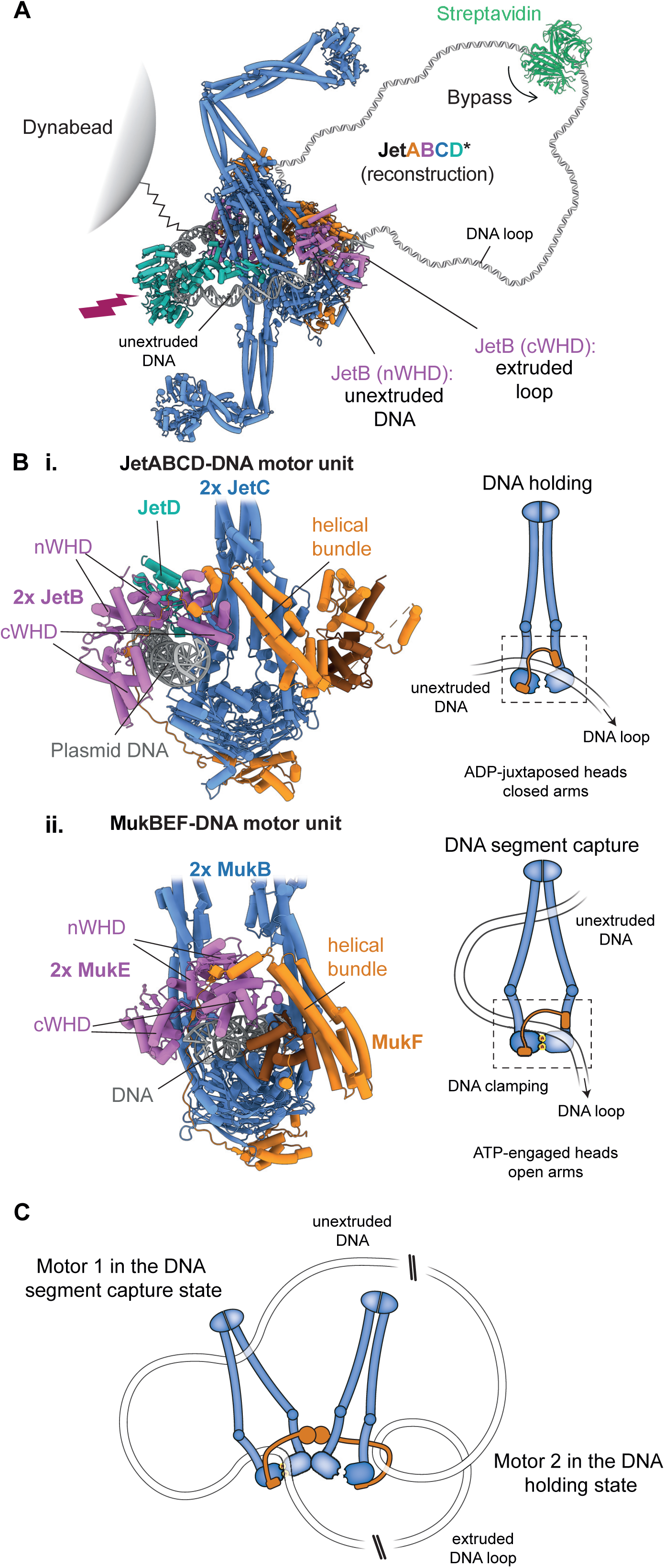
Models for plasmid restriction and loop extrusion. (A) Reconstruction of near full-length, cleavage-competent JetABCD* using AlphaFold2-predicted JetC arms and hinge. Smaller obstacles are bypassed (here represented as a streptavidin tetramer roughly to scale, PDB:1STP ^47^) while larger (micrometer-scaled) impassable obstacles (here as Dynabead) remain unextruded as to-be-cleaved part of the plasmid. The positions of JetB nWHDs and cWHDs are indicated relative to the extruded and unextruded DNA. (B) Alternative states of SMC DNA loop extrusion complexes as represented by structures of plasmid- bound JetABCD* in the cleavage-competent state (i) and *Photorhabdus thracensis* DNA-bound MukBEF (PDB:7NZ0) ^22^ (ii). These conformations correspond to the “DNA holding” and “DNA segment capture” states, respectively ^19,23,32,42^, as schematically illustrated on the right. Note the overall similarity apart from the DNA path and the SMC head and arm conformations. (C) A segment capture model for loop extrusion by JetABCD with one motor unit (right) in the holding and the other (left) in the segment capture state. Note that the motor geometry is as in the resting state. See also Figure S12.

### Cryo-EM analysis of nuclease-defective JetABCD* on extruded plasmid DNA

The DNA visible in the JetABC-DNA structure corresponds to either a small DNA loop captured during initial DNA loading before loop extrusion, or to the final stages of loop extrusion, representing the diminished unextruded segment of plasmid DNA (Figure 3A). To discriminate between these scenarios and to ascertain whether this novel JetABC motor arrangement is indeed primed for plasmid DNA cleavage, we also imaged JetABC in presence of a cleavage-deficient mutant of JetD (E248A, hereafter denoted JetD*) ^8^. We obtained a 3D reconstruction of a plasmid-borne JetABCD* complex at 4.35 Å after performing 3D variability analysis (Figures 3B and S6; Table S1, see Methods). While the architecture of the plasmid-bound JetABC dimer core and its DNA association remained virtually unchanged, the DNA connecting the motor units now exhibited a close association with a JetD* dimer and an altered shape (Figure 3B). The previously continuously bent U-shaped DNA configuration is now transformed into a kinked V-shaped conformation. The structure likely represents a “cleavage-competent” state of the JET nuclease (further discussed below). We further improved the resolution of a JetABCD* core structure to 4.2 Å by enforcing C2 symmetry during non-uniform refinement with a mask (Figures 4A and S6, Table S1, see Methods). This more rigid core includes the JetC heads, the adjacent coiled coils, the JetB dimer bound to DNA, and the JetD aCAP domain bound to the JetC larynx (Figure 4C) ^22^. We then built a model of the plasmid-bound JetABCD* core from this map using a published structure of the resting state (PDB:8AS8 ^8^) and structure predictions as starting material (Figures 4B and S7) ^34–36^.

### Transition from resting to cleavage-competent state

Comparing the model depicting the plasmid-borne, cleavage-competent JetABCD* structure with that of the JetABC resting state reveals multiple similarities and striking differences ^8,9^ (Figures 4A-C and S8A-B). The JetC dimers adopt analogous conformations with heads juxtaposed and arms closed (Figure S8C). Likewise, the JetB dimer remains essentially unchanged, albeit shifted from the side of the JetC heads in the resting state towards their tops (Figures 4B-D and S8A-B). However, the way the kleisin JetA dimer brings together the two JetC motor units is clearly altered (Figure S8D-J). The alterations originate from the folding and interactions of the amino-terminal sequences of kleisin JetA (“JetA-N”): changing both its dimerization (Figure S8D-G) as well as its association with the JetC head-proximal arm, known as the “neck” (Figure S8H-I). JetA-N harbors an amino-terminal winged-helix domain (“nWHD”) followed by a helical bundle (Figures 4B-C and S8A-B,D). Together these domains form a compact dimer in the resting state that connects the two motor units into a V-shaped geometry ^8,9^. In the cleavage-competent state, the JetA-N dimer is more open resulting in an almost I-shaped motor unit configuration (Figures 4A-B and S8A-B). While the nWHD dimer mostly retains its shape (Figure S8F), the helical bundles dissociate and reposition in the cleavage-competent state (moving from the top to the sides of the nWHD dimer; Figure S8D-E). The repositioned helical bundle associates with the JetC neck (Figure S8I). Concomitantly, a segment of the helical bundle—unstructured in the resting state—folds into an alpha helix that also contacts the JetC arm (Figure S8G). Both the substantial DNA bending and the unfolding of the compact JetA-N dimer are potentially energetically unfavorable changes. The driving force behind these structural changes is not immediately apparent. To explore whether the cleavage-competent state emerges upon contact with plasmid DNA also in solution, we performed *in vitro* site-specific cross-linking of JetA and JetC proteins using engineered cysteine residues (Figures 4E and S10). Based on the available structural models, we selected a residue in JetA, namely E229, that is in close proximity to JetC(L267) in the resting state (Cα-Cα 5.9 Å), and to JetC(E840) in the cleavage-competent state (Cα-Cα 7.8 Å) (Figure 4E). Substitution of these residues to cysteines did not markedly impede JetABCD activity *in vitro* (Figure S10A). Furthermore, the JetA subunit was fused with an amino-terminal HaloTag to allow quantification of cross-linked products by in-gel fluorescence detection. Employing bismaleimidoethane (BMOE) cross-linking, we readily detected the JetA-JetC contact in the resting state using the designated reporter cysteines. In contrast, cross-linking involving reporter cysteines for the contact in the cleavage-competent state yielded minimal signal, if any, unless ATP and plasmid DNA were added (Figure 4E). Consistent with the cryo-EM analysis, addition of JetD was neither required nor detrimental for the detection of the JetA-JetC contact in the cleavage-competent state (Figure S10B). We conclude that this contact is indeed formed during JetABCD action on plasmid DNA. Whether the contact is formed exclusively in the cleavage-competent state or also during initial stages of DNA sensing and extrusion remains to be established.

### The DNA-bound JetD nuclease dimer in the cleavage-competent state

Finally, we also produced a local map of the JetD* dimer in the JetABCD*-DNA complex with improved interpretability—mainly within the Toprim domain—by local refinement after local 3D variability analysis around JetD (Figures 5A, S6 and S11A-B; Methods). We observed two main types of architectures, one with a JetD dimer symmetrically attaching to the two κ-JetC proteins—that we describe in more detail here—and another less-well-resolved asymmetric JetD map (Figures S6 and S11). In the two architectures (for symmetric JetD see EMDB:18208 and EMDB:18209), JetABC does not noticeably deviate from the core structure described above (EMDB:18201). We then placed two aCAP domains and a Toprim domain dimer (obtained from AlphaFold2) by rigid-body fitting into the symmetric map of plasmid-bound JetABCD* (Figure 5A). We also incorporated segments of DNA by flexible fitting into the map. While DNA is surrounded by proteins, some parts are exposed to the surface and may thus accommodate the roadblock anchors described above. The central cavity may harbor another DNA double helix, explaining cleavage of catenated DNA circles.

We observe that the JetD* dimer encompasses the entire length of the DNA connecting the motor units and establishes attachment through the aCAP domains to the larynx of the JetC heads (specifically, to the two κ-JetC heads which are bound to carboxy-terminal sequences of kleisin JetA) ^37^ (Figures 3B, 4A, 5A). This interaction is facilitated by a hydrophobic pocket present in JetD aCAP, where the JetC residue F919 inserts (Figures 5A-B and S9B). Notably, perturbing this interface by mutating JetC(F919) or key residues in the JetD pocket to glutamate renders the JET nuclease inactive *in vivo* (Figures 5B and S9C), supporting its physiological relevance. Both the aCAP and Toprim domains align with the DNA, wrapping around the DNA double helix. In contrast to the continuous bending of DNA observed in the absence of JetD*, the DNA in this configuration exhibits a pronounced kink at its midpoint. This kink closely corresponds to where the putative nuclease active site is located on the Toprim domains ^9,10^ (Figures 5A, C). The DNA double helix putatively undergoes this deformation through the combined efforts exerted by the JetABC DNA motors and the JetD nuclease. The deformation may prepare DNA for fitting into the nuclease active site prior to cleavage. The putative active site residues on the pair of Toprim domains are positioned at a small distance apart (JetD E248 Cα-Cα ∼30 Å), likely explaining the short overhangs (1-4 bs) observed in JET-cleaved DNA ^8^. The variability in overhang lengths is possibly explained by DNA flexibility.

We next superposed the JetD* dimer model derived from fitting into the locally refined map (described above, Figure 5A) with available structures of JetD obtained for two other organisms ^9,10^ (Figure 5D). The geometry of the JetD* dimer in plasmid-borne JetABCD* was markedly different from the “closed”, likely autoinhibited, form of an isolated JetD dimer ^9^. In the published “open” conformation ^10^, the aCAP domains are also angled slightly more openly, when compared to the cleavage-competent state (Figure 5D). This is either attributed to different protein origin or, more interestingly, the interaction of the JetD dimer with the JetABC motor units in the cleavage-competent state, possibly with the JetD dimer angle being determined by the aCAP domain binding to the JetC larynx. The geometry of the JetD dimer may also be influenced by the mechanically deformed DNA (Figures 5A, C-D). Notably, rather than passing on top of the Toprim domains and between the aCAP domains of a JetD dimer as proposed before ^9,10^, our structure illustrates that the DNA aligns itself along the V-shaped JetD dimer with the extended shared surface stabilizing the contact. This alignment requires considerable DNA deformations including DNA kinking at the mid-point (Figures 5A, C) likely facilitating DNA access to active site residues. It remains to be elucidated whether such DNA deformations are also needed for accessing the active site of other JET nucleases and the more distantly related Toprim-containing enzymes topoisomerase VI and Spo11 ^27,38,39^ (Figure S11C). AFM images of Spo11-DNA particles and modelling of dimeric complexes from a cryo-EM structure of a monomeric core complex are indeed consistent with the notion of DNA bending ^38,40^. Intriguingly, in Top2, a yeast DNA topoisomerase II (which harbor a Toprim but no arm domain), the G-segment DNA is also strongly bent with the help of Toprim domains (although DNA curvature relative to the Toprim domains is distinct from the one observed in JET) ^41^. DNA bending could thus be a more general feature for the control of Toprim- containing enzymes.

## Discussion

Here we discuss insights into how a bacterial defense system recognizes and neutralizes potentially harmful DNA elements using loop extrusion, a process that otherwise contributes to chromosome folding. We present a model for plasmid restriction by JET nuclease, based on the near-complete extrusion of plasmid DNA followed by bending and cleavage of the residual unextruded segment (depicted in Figure 6A). In addition, we reveal the directionality of SMC translocation, the structure of a DNA holding state as well as blockage of translocation by DNA roadblocks putatively through topological DNA entrapment. Collectively, these findings provide robust support for a DNA segment capture-type model for SMC translocation and loop extrusion ^42^.

### The JET nuclease

The JET system directly recognizes the (pseudo-) circular nature of DNA, probably achieved through the coordinated translocation of oppositely oriented DNA motors along the DNA double helix. This translocation process ultimately extrudes nearly all the DNA culminating in the motors stalling against each other and triggering a transition into the cleavage-competent state. JetD recognizes this cleavage- competent state through direct interactions with the I-shaped motor dimer and with the unextruded U-shaped DNA. The reliance on these two features, both exclusively generated by motor stalling, serves to ensure nuclease activation exclusively on plasmid DNA. However, we note that we cannot rule out that the I-shaped motor dimer might already form earlier, during DNA extrusion, and therefore potentially even on non-plasmid DNA. In principle, encounters between two JET complexes loaded onto a given DNA molecule could also lead to the stalling of motor units. However, such encounters are unlikely to generate the same DNA-motor geometry and are thus expected to fail to activate the JET nuclease function.

MukBEF also forms an I-shaped motor dimer with the helical bundle of kleisin associating with the SMC subunit as here observed for the cleavage-competent state of JET ^8,22^ (Figure S8J). However, in MukBEF, the kleisin helical bundle together with the nWHD display a compact fold, bearing resemblance to the JET resting state. This implies that the dimer geometry of MukBEF contains features of both the JET resting and cleavage-competent states. A configuration with yet another variation in kleisin-SMC contacts (denoted as unclamped state) has been observed in JET when bound to a short linear DNA molecule ^9^. This DNA is localized on ATP-engaged heads, but the KITE dimer is not clamping the DNA and the kleisin N-gate seems open—disconnected from the SMC neck (Figure S12). The role of such an alternative configuration of JET remains to be determined; it could be related to DNA loading or translocation.

When a motor unit encounters a roadblock, it stalls until the second motor unit arrives at the same roadblock position to generate a cleavage-competent state together with the roadblock anchor. The cleavage of the diminishing unextruded segment of plasmid DNA provides an explanation for the observation that roadblocks hinder DNA extrusion yet do not obstruct DNA cleavage, as long as the roadblock can be accommodated within the cleavage-competent state (Figure 6A). This feature may be biologically relevant for example in helping to prevent the emergence of JET resistance by plasmids featuring DNA roadblocks.

### SMC DNA loop extrusion

We present here the structures of the JET nuclease bound to plasmid DNA, unveiling a number of striking features, one of which is the trapped, tight DNA loop connecting two motor units. Although it cannot be ruled out that this configuration is a byproduct of an intricate DNA loading reaction, the fact that the JetD nuclease subunit binds directly to this DNA loop supports the notion that these structures represent a “cleavage-competent” state. This state in all likelihood emerges following circular DNA scanning facilitated by loop extrusion. Accordingly, we can infer the direction of SMC DNA translocation from this structure directly. Taking the JetB KITE dimer as a reference point, we deduce that the DNA emerging from the JetB nWHD dimer corresponds to the “upstream” unextruded DNA, while the cWHD dimer connects to the “DNA loop” representing most of plasmid DNA that has already been extruded (Figure 6). This corresponds to what has been proposed for the segment capture model and also for the related hold-and-feed model when translating from the KITE to the HAWK subunits which support DNA clamping in cohesin and condensin ^23,42^. Curiously, this conclusion contradicts what has been inferred from a cohesin-CTCF-DNA co-structure established on a short linear DNA substrate^43^.

Given the substantial and likely variably-sized SMC translocation steps, it is somewhat unexpected that the DNA U turn linking the two SMC motor units—presumably generated through loop extrusion— exhibits a rather consistent length which enables particle averaging. Our presumption is that the size of this U turn is influenced more by mechanical constraints that accumulate during the final stages of loop extrusion, rather than the sum of all translocation steps culminating in extrusion completion.

The question of whether SMC complexes can bypass large DNA roadblocks is disputed. Some SMC complexes (such as cohesin and condensin) have been reported to exhibit relatively efficient bypass of non-covalently anchored roadblocks ^13,28,43^ in vitro. A more physiological roadblock, a dense Rap1-DNA array, however efficiently blocks condensin translocation with blockage putatively supporting telomere functions *in vivo* ^44^. In our findings, the most plausible explanation is that the JET SMC motor units either cannot bypass or can only inefficiently bypass roadblocks much larger than the dimensions of the complexes themselves. This observation fits well with the cryo-EM structures that indicate topological DNA entrapment, as continuous DNA entrapment would hinder the bypass of large roadblocks. It remains to be fully ascertained whether other SMC complexes, especially those featuring HAWK and larger kleisin subunits, have evolved improved abilities to navigate such roadblocks. However, evidence for stalling of KITE- as well as HAWK-SMC complexes by roadblocks on chromosomes has been reported ^45,46^. Transient DNA leakage with non-covalent anchoring thus remains a valid alternative explanation for the roadblock bypass observed in single-molecule imaging experiments at this juncture.

Whether the dimer geometry observed in the cleavage-competent state is compatible with DNA extrusion is doubtful as the motors are stacked against each other in this state. The DNA clamped by one motor unit were to clash with the other motor unit. Conceivably, an alternative dimer geometry (MukBEF-like or resting state-like) may promote loop extrusion and convert into the cleavage-competent state only upon completion of DNA extrusion. Regardless of dimer geometry, the cleavage-competent state provides key insights into the DNA-motor configuration for a given motor unit. As detailed earlier, the cleavage-competent state harbors motor units with DNA clamped by the JetB KITE dimer onto their SMC heads, held within the kleisin compartment, but not within the SMC compartment ^33^ (Figure 6B). This is distinct from the ATP-bound DNA clamping state—such as observed in the case of MukBEF. Instead, it corresponds to a post-hydrolysis ADP-bound DNA holding state as proposed in the DNA segment capture (and related) model(s) ^19,23,32,42^ while none of the other models for SMC loop extrusion postulated such a state ^1,17,18^. The notion that ADP persists on SMC heads following ATP hydrolysis until the transition into the DNA holding state is complete, as implied by our structures, moreover, provides a rationale for the avoidance of premature ATP engagement. This safeguard mechanism may help prevent unproductive cycles of ATP hydrolysis ^42^. If so, a dedicated mechanism for timely release of ADP may exist. However, we cannot exclude the possibility that ADP accumulating at artificially high levels (through hydrolysis of ATP) during grid preparation led to the ADP occupancy. It will be fascinating to further explore the interconversion of the different states through hydrolysis and binding of ATP.

## Acknowledgments

We are grateful to the Dubochet Centre for Imaging (DCI-Lausanne) for assistance with cryo-EM grid preparation, data collection, and advise for data processing, and to Kyle Muir and Dongchun Ni for support and feedback in data processing. We thank Céline Bouchoux and Masashi Minamino for technical advice. We also thank Bruna de Carvalho and Giorgia Wennubst for technical help in this work. We are grateful to Stéphane Marcand and all members of the Gruber lab for helpful feedback during manuscript preparation.

## Author contributions

F.R.H and H.W.L purified protein, performed biochemistry, and chemical cross-linking experiments.

F.R.H carried out cryo-EM studies. H.W.L performed *in vivo* analyses. M.T. purified MR, provided technical assistance during protein purification and prepared DNA substrates. Y.L. purified JetD(E248A) protein. F.R.H, H.W.L and S.G. wrote the manuscript. S.G. acquired funding and supervised the project.

## Data and materials

Maps and models of the determined structures are available at EMDB and PDB, respectively. All other raw data will be made available via Mendeley Data.

## Methods

### Protein purification

#### Purification and reconstitution of JetABCD

JetABC was purified similarly as previously described ^8^, with the following modifications. After the cell lysate clarification step, the supernatant was loaded onto a 5 mL StrepTrap XT column (Cytiva), followed by 5 column volumes (CV) of washing with lysis buffer (50 mM Tris–HCl pH 7.5, 300 mM NaCl, 5 % (v/v) glycerol, 25 mM imidazole). The complex was eluted with 4 CV of elution buffer (20 mM Tris– HCl pH 8, 200 mM NaCl, 50 mM biotin) and 1 mL fractions were collected. Suitable fractions containing the complex were then pooled and the tag was removed by addition of 3C protease (200 µL of 1 mg/mL) followed by an overnight incubation at 4°C. The resulting solution was concentrated using Amicon Ultracentrifugal filter units (50 kDa cutoff; Millipore) and injected onto a Superose6 Increase 10/300 GL size-exclusion chromatography (SEC) column, either equilibrated with 20 mM Tris–HCl pH 7.5, 250 mM NaCl and 1 mM TCEP, or with ATG buffer (10 mM Hepes–KOH pH 7.5, 150 mM KOAc, 5 mM MgCl_2_) supplemented with 1 mM of TCEP. Output fractions were concentrated to around 15-20 μM and flash-frozen in liquid nitrogen. The complex purified in ATG buffer was used for cryo-EM, streptavidin circles, and cross-linking experiments. JetD was purified as previously described ^8^, except for cross-linking experiments where the SEC step was performed in ATG buffer.

For JetABC complexes harboring Halo-tagged JetA for use in cross-linking assays, the same protocol was followed but the 3C protease cleavage step was omitted. ATG buffer supplemented with TCEP (1 mM final) was used for the final SEC purification step.

JetABCD was reconstituted as previously described ^8^ by mixing JetABC and JetD in ATG buffer (10 mM Hepes–KOH pH 7.5, 150 mM KOAc, 5 mM MgCl_2_) or MM buffer (25 mM Hepes pH 7.5, 250 mM potassium glutamate, 10 mM magnesium acetate) at the appropriate concentration (typically 250 nM of JetABC d-o-p with 1000 nM of JetD).

#### Purification of Mre11-Rad50

Mre11-Rad50 was expressed in *E. coli* BL21 cells from two expression vectors (N-terminal 10His- TwinStrep-3C tag for Mre11, no tag for Rad50). One liter of culture in TB medium was grown at 37°C until OD_600_=1. The culture was then cooled down to 18°C and protein overexpression was induced by IPTG addition (0.5 mM final) for 16 hours. Cells were harvested by centrifugation, resuspended in lysis buffer (50 mM Tris pH7.5, 300 mM NaCl, 5% (v/v) glycerol, 25 mM imidazole) freshly supplemented with 100 μL of protease inhibitor cocktail (Sigma P8849), and 5 mM 2-mercaptoethanol. Cells were lysed by sonication on ice using a with a VS70T probe mounted on a SonoPuls unit (Bandelin), at 40% output for 13 min with pulsing (1 s on / 1 s off). After clarification by ultracentrifugation (40,000 g for 45 min), the lysate was loaded onto an HisTrap HP 5 mL column (Cytiva). After 5 CV (column volume) washes in lysis buffer, proteins were eluted by a gradient elution in lysis buffer containing imidazole (25-500 mM). Fractions containing the complex were then pooled, diluted in 20 mM Tris pH 7.5, 50 mM NaCl, and loaded onto a 5 mL HiTrap Q column (Cytiva). After 5 CV washes in 20 mM Tris pH 7.5, 50 mM NaCl, proteins were eluted by an increasing NaCl gradient (50-1000 mM). Mre11 and Rad50 containing fractions were pooled and concentrated using Amicon Ultracentrifugal filter units (50 kDa cutoff; Millipore) and injected onto a Superose6 Increase 10/300 GL SEC column (equilibrated with 20 mM Tris–HCl pH 7.5, 250 mM NaCl and 1 mM TCEP). Final complex-containing fractions were pooled, concentrated again to about 0.9 mg/mL and flash frozen in liquid nitrogen for long term storage at - 70°C.

#### Purification of SNAP-tag

SNAP-tag was produced in *E. coli* BL21, expressed with a carboxy-terminal 3C-8His tag. A one-liter culture of the BL21 strain was grown in TB medium at 37°C until the culture reached OD_600_=0.5. The culture was cooled to 16°C and protein overexpression induced by IPTG (0.5 mM final) for 16 hours. Cells were harvested by centrifugation, resuspended in lysis buffer (50 mM Tris pH 7.5, 300 mM NaCl, 5% (v/v) glycerol, 25 mM imidazole), followed by lysis via sonication on ice with a VS70T tip using a SonoPuls unit (Bandelin), at 40% output for 12 min with pulsing (1 s on / 1 s off). After clarification by ultracentrifugation (40,000 g for 45 min), the lysate was loaded onto a HisTrap HP 5 mL column (Cytiva), washed with 6 CV of lysis buffer and eluted with lysis buffer containing an increasing imidazole gradient (up to 500 mM) for 10 CV. Fractions containing SNAP-3C-8His were collected and dialyzed back to lysis buffer containing 3C protease overnight at 4°C. To remove the His-containing cleaved epitope and 3C protease, the dialyzed sample was loaded onto a HisTrap column and washed with 5 CV lysis buffer, the SNAP-containing flowthrough was collected. SNAP was then dialyzed into PBS buffer for use in DNA cleavage assays. SNAP-tag coupling activity (methodology described below) was first tested with a small ligand BG-biotin (NEB) or DNA (a 550 bp benzylguanylated PCR product) by SDS-PAGE and agarose gel analysis respectively (Figures S4F-G). We note that SNAP-DNA coupling is a much less efficient reaction than with small ligands, requiring a molar excess of SNAP to achieve high coupling rates.

### DNA substrate preparation

#### Artificial DNA circles

The 2.3 kb end-biotinylated DNA substrates were produced by PCR amplification from *B. subtilis* genomic DNA using STO331/STG995 (double-biotinylated) and STO337/STG995 (single-biotinylated) primers pairs (see Table S2). Excess primers were removed by column purification. The DNA was then phenol-chloroform extracted, followed by salt/ethanol precipitation and solution in water. Artificial DNA circles were generated by streptavidin addition to the 2.3 kb DNA (150 ng per 15 µL reaction, corresponding to 6.7 nM) diluted in ATG buffer supplemented by 1 mM of MnCl_2_. For specific enrichment of circular species (for Figure S1C), DNA species obtained after streptavidin addition were subjected to Rad50/Mre11 treatment (125 nM final tetramer) for 5 minutes at 37°C, followed by inactivation by incubation 5 min at 55°C. We note variable efficiency of pseudo-circle generation. The actual final concentration of streptavidin (50-100 nM monomer) was selected based on the enrichment of DNA species generated prior to experimentation (circular and linear, see Figures 1B and S1B-C).

#### Chemically modified DNA circles

See Table S2 for a list of modified substrates and their starting plasmid/oligo(s). Modified DNA substrates (2.9 kb circles containing ssDNA flaps, biotin and benzylguanine) were prepared via a gap- replacement approach described in^30^. 4-5 µg of pSG6970/pG46^55^ or pSG7084 (containing one (with site A) and two (with sites A and B) oligo replacement cassettes respectively, see Figure S2A for schematic) was nicked in rCutsmart buffer (NEB) with Nt.BbvCI (NEB, 30 U) in 20 µL reactions for one hour at 37°C. The reaction was quenched by addition of 3 µL Tris pH 8 (200 mM) and 3 µL EDTA pH 8 (400 mM). An excess of replacement oligo (4 µL of 100 µM stock) was added to the mix. Nt.BbvCI-nicked ssDNA fragments were melted by heating at 80°C for two minutes, followed by oligo annealing by slowly cooling the reaction to 20°C (1°C decrease every minute). After column purification and elution in 52 µL water, DNA nicks were sealed by addition of 6 µL T4 ligase buffer (Thermo Fisher) and 2 µL T4 ligase (10 U, Thermo Fisher), followed by incubation for one hour at 22°C. Finally, a second round of column purification was performed.

For doubly modified substrates, the replacement reaction was performed by adding two replacement oligos into the tube containing nicked pSG7084.

For DNA circles containing ssDNA gaps, a variation of the protocol was used: pSG6969/pG68 ^55^ (825 ng) was nicked with Nb.BbvCI (NEB, 10 U) in 25 µL reactions for one hour at 37°C. To capture the nicked ssDNA fragments, an excess of complementary oligo mix (2.5 µL of 5 µM, see Table S2) was added, followed by heating at 80°C for two minutes, and oligo annealing by slowly cooling the reaction to 20°C (1°C decrease every 15 s). The annealed short dsDNA fragments were then removed by column purification, retaining circles containing ssDNA gaps.

#### Catenated DNA

DNA catenanes were prepared as previously described ^31^. pSG6971 (50 µg) was incubated with Tn3 resolvase (2 µM final) in Tn3 buffer (50 mM Tris pH8, 70 mM MgCl_2_, 0.1 mM EDTA) in 200 µL reactions at 37°C for 4 hours, followed by heat inactivation at 65°C for 20 minutes. DNA was then purified via phenol:chloroform extraction.

### DNA cleavage assay

DNA cleavage reactions were performed as described ^8^. Unless stated otherwise, DNA was incubated with JET (12.5 nM d-o-p) in ATG buffer (10 mM Hepes-KOH pH 7.5, 150 mM KOAc, 5 mM MgCl_2_) supplemented with 1 mM ATP in 15 µL reactions, at 37°C for 15 minutes. For reactions involving restriction enzymes, 5 U was typically used under the same reaction conditions. For flapped, gapped and catenated DNA substrates, 3.5 nM, 4.5 nM, 8.1 nM of plasmid DNA were used, respectively. For experiments involving artificial DNA circles made by streptavidin addition to biotinylated DNA, 1 mM of MnCl_2_ was added to the ATG buffer. Addition of MnCl_2_ does not impact JET activity. After artificial circle formation described above, DNA was subjected to JET (12.5 nM d-o-p), RecBCD (NEB, 5 U), NcoI-HF (NEB, 5 U) or Rad50/Mre11 (125 nM final tetramer) treatment for 15 min at 37°C. The resulting DNA species were resolved in 1% agarose (without SDS) gels containing EtBr.

### Experiments with DNA substrates anchored to streptavidin-coated Dynabeads

#### Cleavage assay

(Per reaction, scaled up accordingly) M-270 streptavidin-coated Dynabeads (10 µL containing 100 µg, Thermo Fisher) were equilibrated and washed with 1xBW buffer (5 mM Tris-HCl pH 7.5, 0.5 mM EDTA, 1 M NaCl) according to manufacturer’s protocol. The beads were resuspended in 20 µL 2xBW buffer (10 mM Tris-HCl pH 7.5, 1 mM EDTA, 2 M NaCl) and incubated with an equal volume of biotinylated DNA (500 ng in water) at 30°C for 30 minutes with shaking. After washing with 1xBW buffer, the beads were equilibrated with MM reaction buffer (25 mM Hepes pH 7.5, 250 mM potassium glutamate, 10 mM magnesium acetate, 1 mM DTT).

The beads were then resuspended in MM buffer supplemented with 1 mM ATP (15 µL per reaction). The control unmodified DNA (50 ng per reaction) in this buffer (15 µL per reaction) was directly added into Protein G dynabeads (Thermo Fisher) that were washed and equilibrated as above. For experiments involving multiple conditions, this was split into 15 µL aliquots. Beads were treated with JET (12.5 nM d-o-p) and/or ScaI-HF (NEB, 5 U) for 15 minutes at 37°C. DNA bound to the beads was eluted by addition of 85 µL preheated MM buffer supplemented with 25 mM biotin and SDS (0.1% final) and incubation for 10 minutes at 70°C. The supernatant containing the eluate was separated from the beads with a magnetic rack. To remove salt that hindered DNA migration and visualization in agarose gels, the eluted DNA was column purified into 15 µL water. This was followed by addition of SDS (0.5% w/v final) containing loading buffer. The reactions were loaded onto a 0.03 % (w/v) SDS- and EtBr-containing 1 % (w/v) agarose gel, ran at 5V/cm for 1 hour and bands visualized with on a transilluminator (UVP GelSolo). For the experiment with linear biotinylated DNA (same as the ones used for the artificial circles), the same procedure was followed but with inclusion of MnCl_2_ (1 mM final) in MM buffer and Rad50/Mre11 tetramer (125 nM final).

#### DNA leakage assay

For the DNA leakage assay, the same protocol was followed but without addition of enzymes. After mock treatment of beads in MM buffer with or without 25 mM biotin for 15 minutes at 37°C, the supernatant containing the released fraction was isolated. The beads containing still-bound material was eluted by the addition of 15 µL preheated MM buffer supplemented with 25 mM biotin and SDS (0.1% final) for 10 minutes at 70°C. The supernatant containing the elution fraction was collected from the beads. Both fractions were column purified and loaded onto an EtBr gel. To quantify DNA leakage from beads, the intensity of bands (obtained from ImageJ) of the released fraction was divided from the sum of the released and elution fractions.

### Experiments with DNA substrates covalently linked to dynabeads

#### Coupling BG-linked ligands to SNAP

In 25 µL reactions, BG-labelled substrate (150 nM 550 bp PCR product, 40 nM 2.9 kb DNA circles or 1- 50 µM BG-biotin) in PBS was added to purified SNAP (2.5 or 5 µM final) in the presence of 1 mM DTT for 2 hours (or as indicated) at 37°C. If the coupled substrate is to be reacted with dynabeads, 75 µL of PBS was then added to the reaction, resulting in 100 µL of coupled DNA.

#### Coupling SNAP-DNA with Epoxy Dynabeads

(Per coupling reaction) 4 mg of M-270 Epoxy Dynabeads (Thermo Fisher) were equilibrated and washed in Epoxy A buffer (0.1 M sodium phosphate). After resuspension in 100 µL Epoxy A buffer, 100 µL of SNAP-DNA was added, followed by addition of 100 µL Epoxy B buffer (3 M (NH_4_)_2_SO_4_). The mixture was incubated at 37°C for 16 hours, under slow rotation. To remove uncoupled material and to quench any free epoxy groups, the beads were then rigorously washed with PBS supplemented with BSA (0.1% w/v final), followed by washes in ATG buffer supplemented with BSA (0.1% w/v final).

To confirm that the washed beads were coupled with DNA circles through SNAP, beads (0.5 mg per condition) were treated with either ScaI+EcoRI (5 U each) or proteinase K (1 µL of 20 mg/mL stock, Eurobio) for 30 minutes at 37°C, this resulted in release of band(s) of the expected sizes to the supernatant (Figure S4H). No DNA was coupled in SNAP-tag absence (Figure S4H).

#### Cleavage assay

For each experimental condition, 0.5 mg coupled beads were used. Washed beads were resuspended in ATG buffer supplemented with 1 mM ATP (15 µL per reaction). The control unmodified DNA (50 ng per reaction) in this buffer (15 µL per reaction) was directly added into epoxy beads (0.5 mg per reaction), washed and equilibrated as above. Beads were split into 15 µL aliquots per condition and were treated with JET (12.5 nM d-o-p) and/or ScaI-HF (5 U) for 15 minutes at 37°C. Elution was carried out by addition of 1 µL proteinase K (20 mg/mL, Eurobio) followed by incubation at 55°C for 30 minutes. The eluate was separated from the beads, column purified into 15 µL water, mixed with of SDS (0.5% w/v final)-containing loading buffer. The reactions were loaded onto a 0.03 % (w/v) SDS- and EtBr-containing 1 % (w/v) agarose gel, ran at 5 V/cm for 1 hour and bands visualized with on a transilluminator (UVP GelSolo). To quantify cleavage efficiency, the intensity of the band of the linear DNA was divided by the sum of all band intensities with ImageJ.

### *in vitro* cysteine cross-linking

Cysteine cross-linking was adapted from ^19^. Reactions were conducted in darkness in ATG buffer supplemented with TCEP (1 mM final). To label Halo-tagged JetA, 1 µL of HaloTag TMR ligand (Promega, 5 mM) was added to 100 µL of JetABC (2.5 µM d-o-p) and incubated for 30 minutes at 37°C. Protein was eluted away from unreacted ligand via Zeba™ Spin Desalting Columns (Thermo Fisher). TMR-labelled JetABC complexes (0.5 µM d-o-p final) were mixed with/without JetD (1 µM dimer final), ATP/ATPγS (1 mM final) and DNA (1.8 kb circular pDonor, 100 nM final) for 10 minutes at room temperature. 0.5 µL of BMOE (20 mM stock) was then added to the mix for 15 seconds before quenching by addition of 1 µL DTT (100 mM stock). The cross-linked reactions were analyzed via SDS-PAGE followed by imaging JetA-TMR with an Amersham Typhoon™ laser scanner (Cy3 settings, PMT auto) and CBB staining.

### Strain construction in *E. coli*

*E. coli* K12 strains containing GF4-3 *jetABCD*, under the arabinose-inducible P_BAD_ promoter and integrated into the neutral chromosomal *glmS* loci, were derived via tri-parental mating as previously described ^56^. The test plasmid pBAD was introduced to electrocompetent cells via electroporation at 2.0 kV. See Table S4 for a list of bacterial strains used.

### Plasmid stability assay in *E. coli*

Plasmid stability assays were performed as described ^8^. Plasmid (pBADMycHisA, short pBAD) and P_BAD_*- jetABCD*-containing *E. coli* cultures were grown overnight in LB supplemented with ampicillin at 37°C. They were diluted into LB at an OD_600_ = 0.0025, with or without arabinose (0.2% (w/v) final) and allowed to grow for approximately ten generations. 200 µL of culture was harvested and serially diluted sevenfold (from 10^-1^ to 10^-7^) in PBS. For total cell number count, 5 µL of the dilutions from 10^-4^ to 10^-7^ were spotted onto nutrient agar plates in duplicate. For pBAD-containing cells, the same amount of all dilutions was spotted onto nutrient agar plates supplemented with ampicillin (100 µg/mL final). After overnight incubation at 37°C, colonies were counted and extrapolated based on the dilution degree to give an estimate of the total cell number and survivor count. To quantify pBAD+ cells, the percentage of ampicillin-resistant cells over the total cell number was calculated. Datapoints greater than 100 % were sometimes obtained if the number of pBAD+ colony count was greater than total cell colony count.

### Cryo-electron microscopy

#### Sample preparation and data collection

JetABC and JetABCD* (JetD*: JetD(E248A)) complexes (800 nM JetABC d-o-p, 1720 nM JetD d) were freshly reconstituted by dilution in ATG buffer (with 1 mM TCEP) containing 100 nM plasmid DNA (pDonor). Complexes were then supplemented with ATP and β-octyl glucoside (final concentration 1 mM and 0.05 % (w/v), respectively), incubated at room temperature for 10 minutes, and cooled on ice prior to freezing. Cryo-EM grids (Au-flat 1.2/1.3 on 300 gold mesh, Jena Bioscience) were freshly glow discharged in an EasyGlow device with 15 mA current, 90 s glow time and 60 s wait time. Then, 3 µl of JET-DNA samples were applied on the grids mounted in a Vitrobot Mark IV, set to 10°C in the chamber and 100% humidity. The grids were blotted at blot force 10 for 0.5 s and vitrified in liquid ethane precooled to liquid nitrogen temperature. The grid screening and exploratory dataset acquisition were performed on a Glacios Cryo-TEM equipped with Falcon IV CMOS detector (Thermofisher Scientific (TFS)) with 150 000x magnification, resulting in a nominal pixel of 0.92 Å and with a total dose of 40 electrons per square angstrom (e-/Å2). High resolution data collection was performed on a 300 kV Titan Krios equipped with a Falcon IV G4i camera (TFS) in counted mode at a magnification of 96 000x (pixel size of 0.83 Å) and data was saved in the EER format using the EPU software (TFS). For the first dataset (JetABC + plasmid DNA), 13,292 movies were collected at a defocus range from -0.6 to -2.4 µm, with a total dose of 50 e-/Å2. For the second dataset (JetABC + JetD(E248A) + plasmid DNA), 37,902 movies were collected at a defocus range from -0.8 to -2.6 µm with a total dose of 40 e-/Å2.

#### Data processing

Initial data pre-processing was performed on-the-fly using cryoSPARC live (V4.1), with further processing using cryoSPARC (V3.3) ^57^. For the plasmid bound JetABC reconstruction (Figures 3 and S5; Table S1), 12,287 dose-weighted micrographs (out of 13,292) were selected based on CTF fit and ice quality. Particles were initially picked using blob picker, followed by extraction at a box size of 80 pixels (the particles were Fourier-cropped 8 times from an initial box size of 640 pixels) which gave a stack of 1,257,512 particles. After several rounds of 2D classification, the particles were re-extracted (box size of 320 pixels, Fourier-cropped 2 times) and subjected again to 2D classification. The best 224,413 particles were used for an *ab initio* reconstruction with 3 classes. Non-uniform refinement ^58^ (C2 symmetry enforced) of the best class (class 3, 103,418 particles) led to an initial reconstruction that was used to generate 2D templates. 1,492,370 particles were obtained after template picking and particle extraction (box size: 80 pixels, Fourier-crop: 8). After several rounds of 2D classification and particle re-extraction at Fourier-crop 2 (initial extraction box: 640 pixels), the 375,312 selected particles were subjected to an *ab initio* reconstruction with 3 classes followed by one round of heterogenous refinement. Non-uniform refinement (first with C1, then with C2 symmetry enforced) of the best class (class 3, 187,999 particles) gave the reconstruction of the plasmid bound JetABC core at an overall resolution 4.78 Å. The map was finally subjected to local filtering in cryoSPARC.

For the plasmid bound JetABCD* reconstructions (Figures 3, 4, 5, S6, S7 and S11; Table S1), 29,566 dose-weighted micrographs (out of 37,902) were selected based on CTF fit and ice quality. Particle picking using the blob picker followed by particle extraction (box size: 640 pixels, Fourier-cropped 8 times to 80 pixels) gave an initial stack of 2,316,479 particles. After several rounds of 2D classification and re-extraction (box size: 640 pixels, Fourier-cropped two times to 320 pixels), the 176,397 selected particles were subjected to *ab initio* reconstruction with 2 classes. Non-uniform refinement of the best class gave an initial volume (C1 symmetry, 5.82 Å) that was used to create the 2D templates. After template picking and particle extraction, (box size: 640 pixels, Fourier-cropped 8 times to 80 pixels) a new stack of 3,761,906 particles was obtained. After cleanup by several rounds of 2D classification followed by particle re-extraction (box size: 640 pixels, Fourier-cropped 2 times to 320 pixels), the 659,766 selected particles were used for a 4 classes *ab initio* reconstruction followed by heterogenous refinement. Non-uniform refinement (C1 enforced) of the best class gave a consensus reconstruction of the plasmid bound JetABCD* at an overall resolution of 4.5 Å. In this consensus map, the JetABC core was better resolved, while the “unextruded” DNA part bound by JetD* remained less clear. Thus, 3D variability analysis (4 modes, filter resolution 6.6 Å) was performed ^59^ (Figures S6 and S11A). 3D variability display in frames (Fourier-cropped 4 times to a box size of 160 pixels, Figure S11A) revealed motion within the complex. The JetABC core appeared to have intrinsic flexibility, where the JetA bundle-JetC coiled coils interface and the JetA nWHD dimer were the most rigid parts. In contrast, JetD* bound to the “unextruded DNA” region of the complex was found to have more drastic motion, suggesting that JetD* has a higher degree of flexibility and can adopt different conformations. 3D variability display using two clusters was used to sort these conformations. The best-resolved cluster gave the JetABCD* cleavage-competent state reconstruction after non-uniform refinement (first with C1 then with C2 symmetry enforced) at an overall resolution of 4.35 Å. The latter reconstruction was subjected to local filtering. To improve the JetABCD* core, non-uniform refinement using all the consensus particles with C2 symmetry enforced and a mask around the less flexible regions of the complex (JetABC core and the N-terminal region of the JetD* aCAP) gave the reconstruction of JetABCD core at an overall resolution of 4.2 Å. To further improve the interpretability of the JetD*/DNA region, 3D variability analysis with a mask around JetD* (4 modes, filter resolution 10) followed with 3D variability display in 8 clusters was performed. The best resolved class of JetD* (representing the JetD* region of the cleavage-competent state) was further subjected to homogenous refinement followed by a local refinement using a mask around JetD, which gave the reconstruction of the JetD*/DNA region in the cleavage-competent state at an overall resolution of 6.34 Å. As the plasmid bound JetABCD* reconstructions were all obtained in the wrong hand, the hand was flipped for the deposited and depicted material using ChimeraX (except for Figure S6 where the maps are shown as obtained). Of note, while the position of the upper part of JetC coiled coils and hinges are discernable as fuzzy density (not shown in the main figures, but more clearly visible in the *ab initio* volumes (Figures S5, S6), we could not obtain a suitable reconstruction despite attempts of local refinement.

#### Model building and data presentation

For the model building of the JetABCD* core in the cleavage-competent state, we used as starting material our previous high resolution model of the JetABC monomer (PDB:8AS8 ^8^) and Alphafold2 predictions (a precedent JetA N-terminal/JetC head model ^8^ or newly predicted using Alphafold2 (JetC head-JetD aCAP, JetC head-JetA C-terminus and JetA bundle- other JetA helix) run on the UNIL computing cluster ^34,36^. The models were segmented, rigid body docked and flexibly fitted into the JetABCD core map using ChimeraX (V1.4) ^60,61^ and ISOLDE (V1.6) ^62,63^. The plasmid DNA was modelled *de novo* as idealized B-form polyAT tracks and flexibly fitted into the map using Coot (V0.9.8.3) ^64,65^. The model was manually rebuilt into the density and iteratively improved using Coot/ISOLDE and real space refinement in PHENIX (V1.20.1) ^66,67^. The model was validated using MolProbity ^68^ implemented in PHENIX. Chimera ^69^ and ChimeraX ^60,61^ were used for figure preparation.

## Supplementary tables

Table S1: Cryo-EM data collection and statistics.

Table S2: List of oligonucleotides and chemically modified DNA substrates.

Table S3: Plasmid list.

Table S4: Bacterial strain list.

