## Supplementary Figures for "Structural basis for plasmid restriction by SMC JET nuclease"

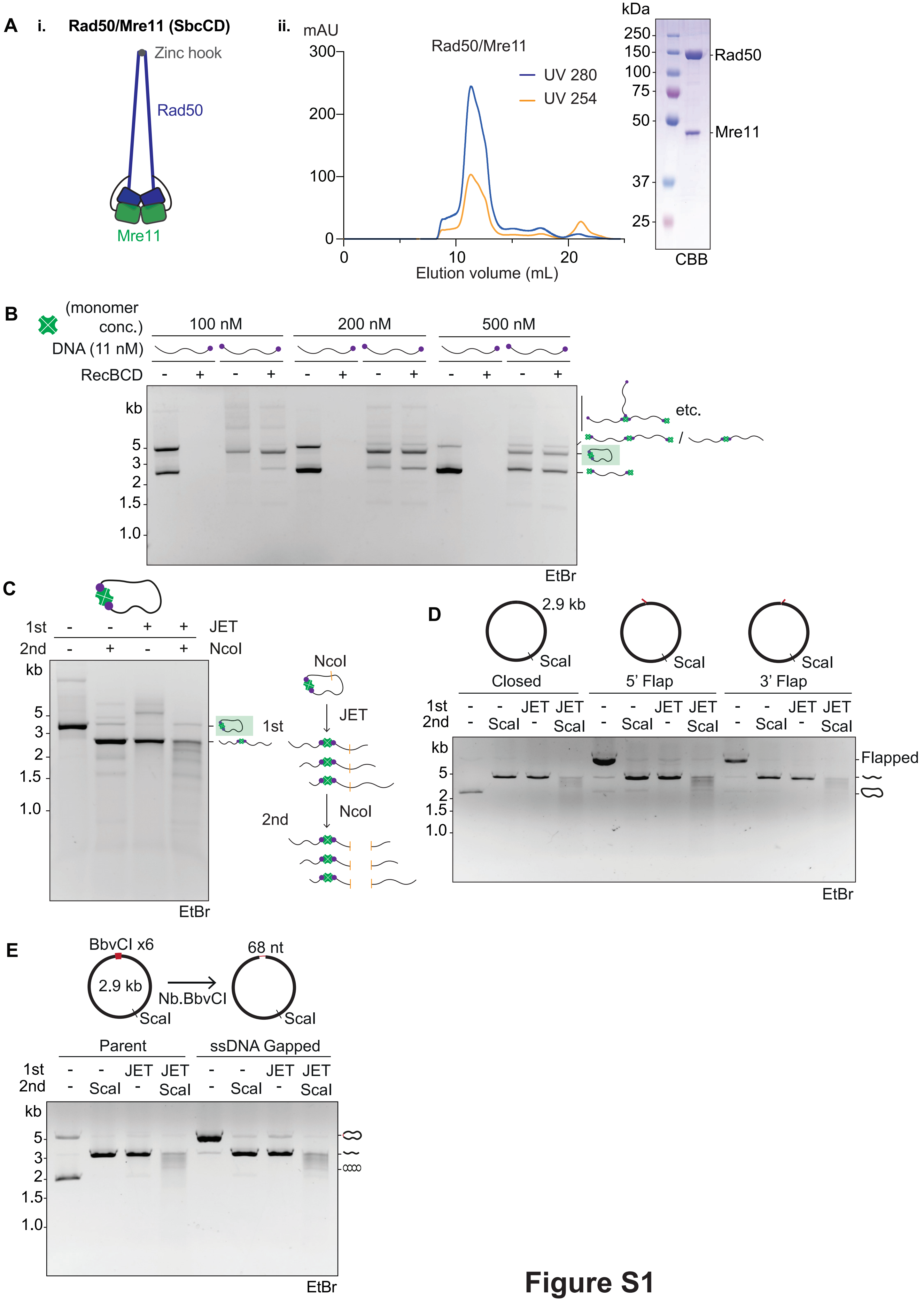

A
i.
Rad50/Mre11 (SbcCD)
Zinc hook
Rad50
Mre11
ii.
mAU
Rad50/Mre11
UV 280
UV 254
Elution volume (mL)
kDa
Rad50
Mre11
CBB
B
(monomer conc.)
100 nM
200 nM
500 nM
DNA (11 nM)
RecBCD
kb
etc.
EtBr
C
1st
2nd
JET
NcoI
kb
EtBr
D
2.9 kb
ScaI
Closed
5' Flap
3' Flap
1st
2nd
JET
ScaI
kb
Flapped
EtBr
E
BbvCI x6
2.9 kb
Nb.BbvCI
68 nt
ScaI
Parent
ssDNA Gapped
1st
2nd
JET
ScaI
kb
EtBr


### Figure S1

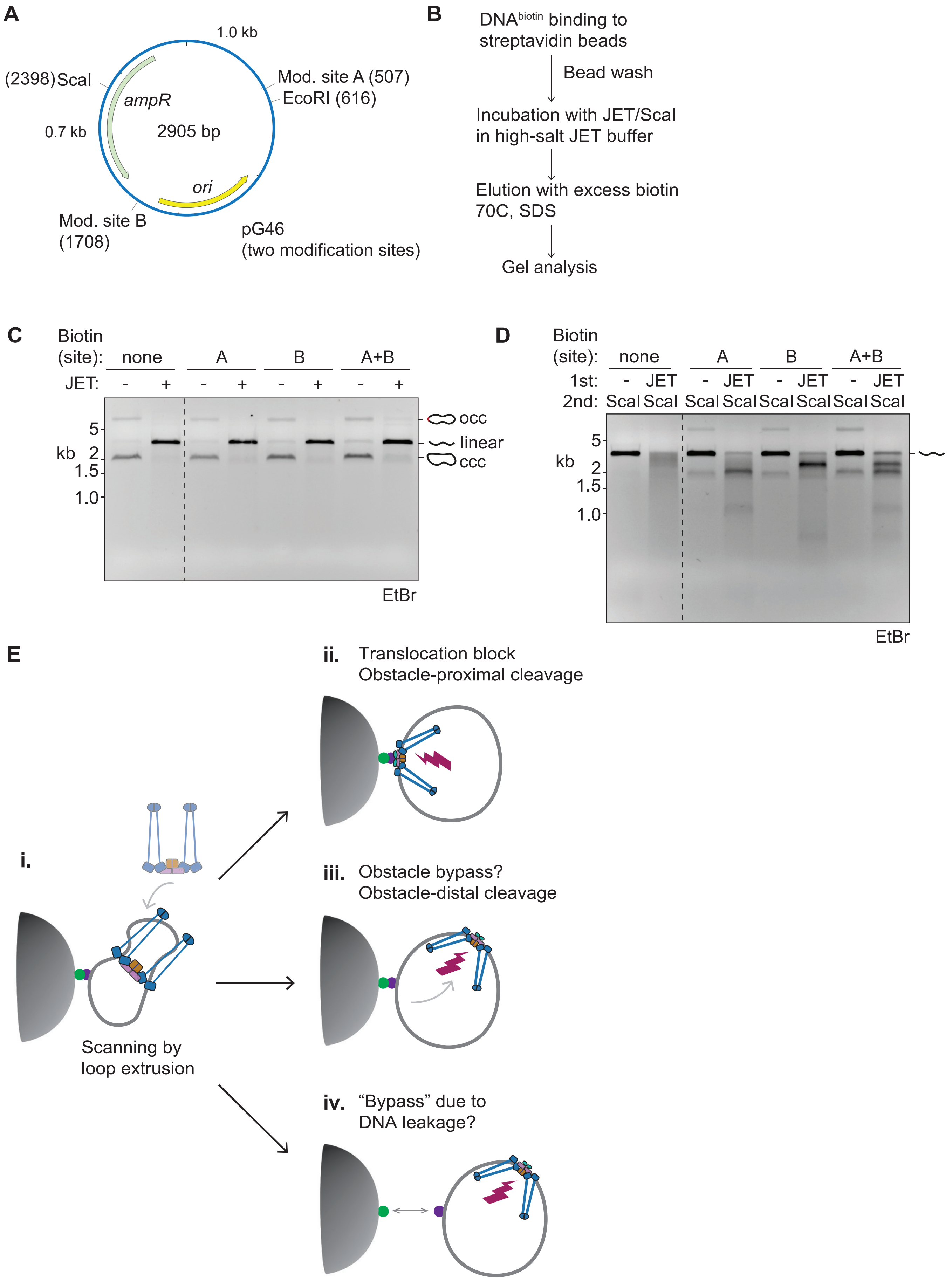
A
1.0 kb
(2398) ScaI
ampR
Mod. site A (507)
EcoRI (616)
0.7 kb
2905 bp
ori
Mod. site B
(1708)
pG46
(two modification sites)
B
DNAbiotin binding to streptavidin beads
Bead wash
Incubation with JET/ScaI in high-salt JET buffer
Elution with excess biotin 70C, SDS
Gel analysis
C
Biotin (site): none A B A+B
JET: - + - + - + - +
kb 5 2 1.5 1.0
occ
linear
ccc
EtBr
D
Biotin (site): none A B A+B
1st: - JET - JET - JET - JET
2nd: ScaI ScaI ScaI ScaI ScaI ScaI ScaI ScaI
kb 5 2 1.5 1.0
EtBr
E
i.
Scanning by loop extrusion
ii. Translocation block
Obstacle-proximal cleavage
iii. Obstacle bypass?
Obstacle-distal cleavage
iv. “Bypass” due to
DNA leakage?

### Figure S2

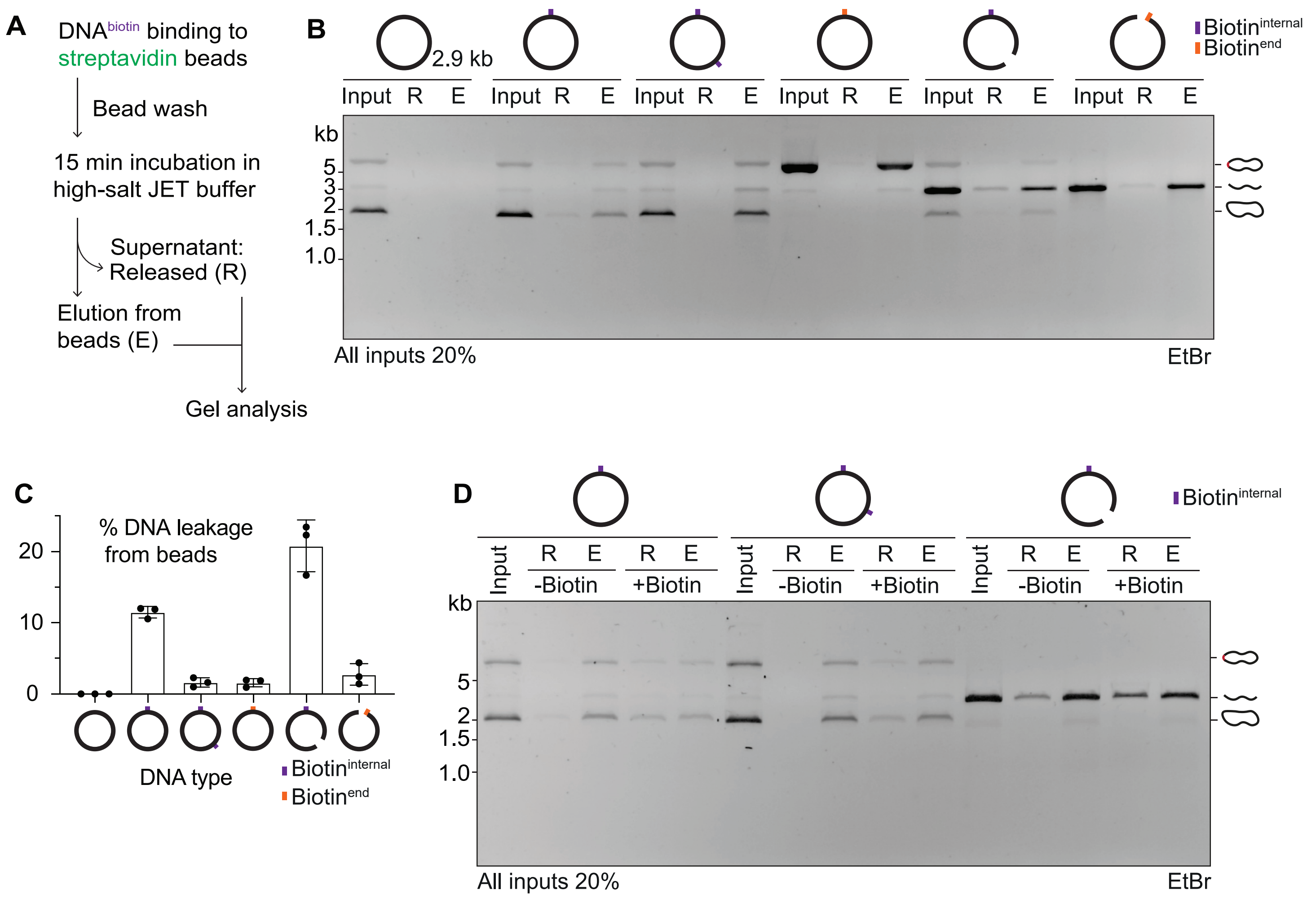
A
DNAbiotin binding to streptavidin beads
Bead wash
15 min incubation in high-salt JET buffer
Supernatant: Released (R)
Elution from beads (E)
Gel analysis
B
2.9 kb
Biotininternal
Biotinend
Input R E
kb
5
3
2
1.5
1.0
All inputs 20%
EtBr
C
% DNA leakage from beads
20
10
0
DNA type
Biotininternal
Biotinend
D
Biotininternal
Input
R E
-Biotin
+Biotin
kb
5
2
1.5
1.0
All inputs 20%
EtBr

**Figure S3**

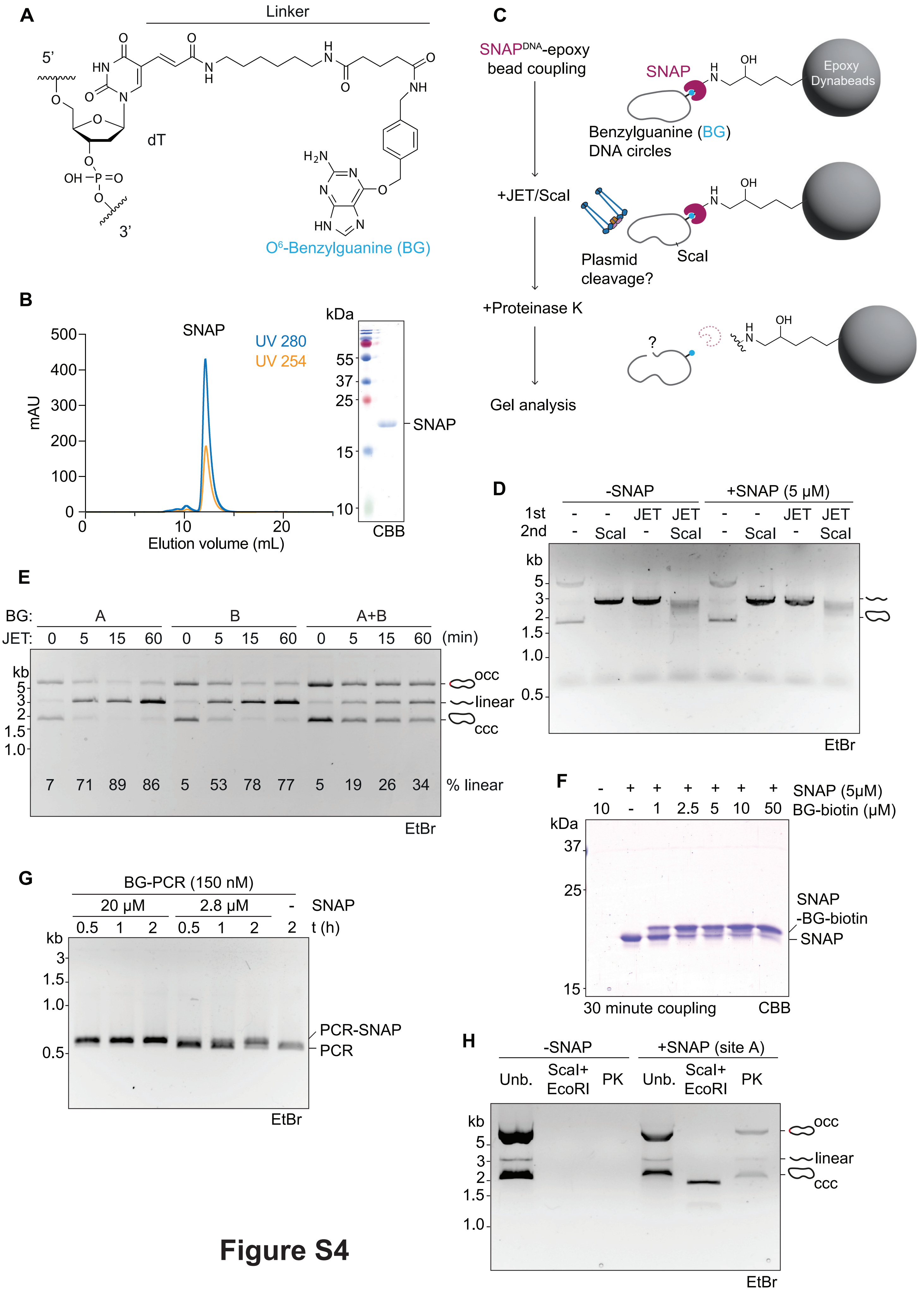
A
Linker
5'
3'
dT
O6-Benzylguanine (BG)
B
SNAP
UV 280
UV 254
mAU
Elution volume (mL)
kDa
55
37
25
15
10
SNAP
CBB
C
SNAPDNA-epoxy bead coupling
SNAP
Epoxy Dynabeads
Benzylguanine (BG) DNA circles
+JET/ScaI
Plasmid cleavage?
ScaI
+Proteinase K
Gel analysis
D
-SNAP
+SNAP (5 µM)
1st
2nd
JET
ScaI
kb
EtBr
E
BG: A B A+B
JET: 0 5 15 60 0 5 15 60 0 5 15 60 (min)
occ
linear
ccc
7 71 89 86 5 53 78 77 5 19 26 34 % linear
EtBr
F
SNAP (5µM)
10 - 1 2.5 5 10 50 BG-biotin (µM)
kDa
SNAP -BG-biotin
SNAP
30 minute coupling
CBB
G
BG-PCR (150 nM)
20 µM 2.8 µM - SNAP
0.5 1 2 0.5 1 2 2 t (h)
PCR-SNAP
PCR
EtBr
H
-SNAP
+SNAP (site A)
Unb.
ScaI+ EcoRI
PK
occ
linear
ccc
EtBr

### Figure S4

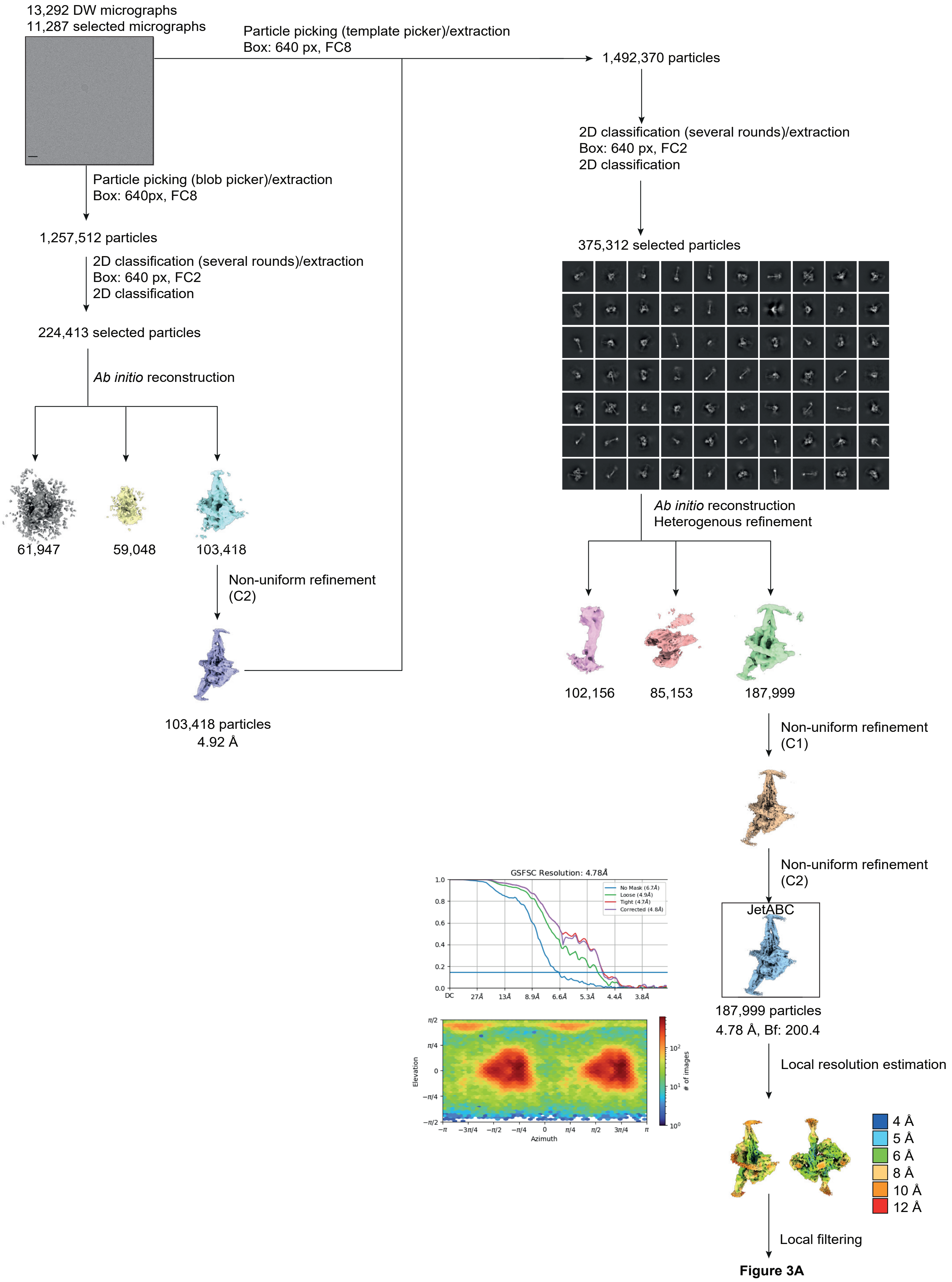

13,292 DW micrographs
11,287 selected micrographs
Particle picking (template picker)/extraction
Box: 640 px, FC8
1,492,370 particles
2D classification (several rounds)/extraction
Box: 640 px, FC2
2D classification
375,312 selected particles
Particle picking (blob picker)/extraction
Box: 640px, FC8
1,257,512 particles
2D classification (several rounds)/extraction
Box: 640 px, FC2
2D classification
224,413 selected particles
Ab initio reconstruction
61,947
59,048
103,418
Non-uniform refinement
(C2)
103,418 particles
4.92 Å
Ab initio reconstruction
Heterogenous refinement
102,156
85,153
187,999
Non-uniform refinement
(C1)
Non-uniform refinement
(C2)
JetABC
187,999 particles
4.78 Å, Bf: 200.4
GSFSC Resolution: 4.78Å
No Mask (6.7Å)
Loose (4.9Å)
Tight (4.7Å)
Corrected (4.8Å)
DC
27Å
13Å
8.9Å
6.6Å
5.3Å
4.4Å
3.8Å
Elevation
Azimuth
### of images
Local resolution estimation
4 Å
5 Å
6 Å
8 Å
10 Å
12 Å
Local filtering
Figure 3A


### Figure S5

### Figure S6

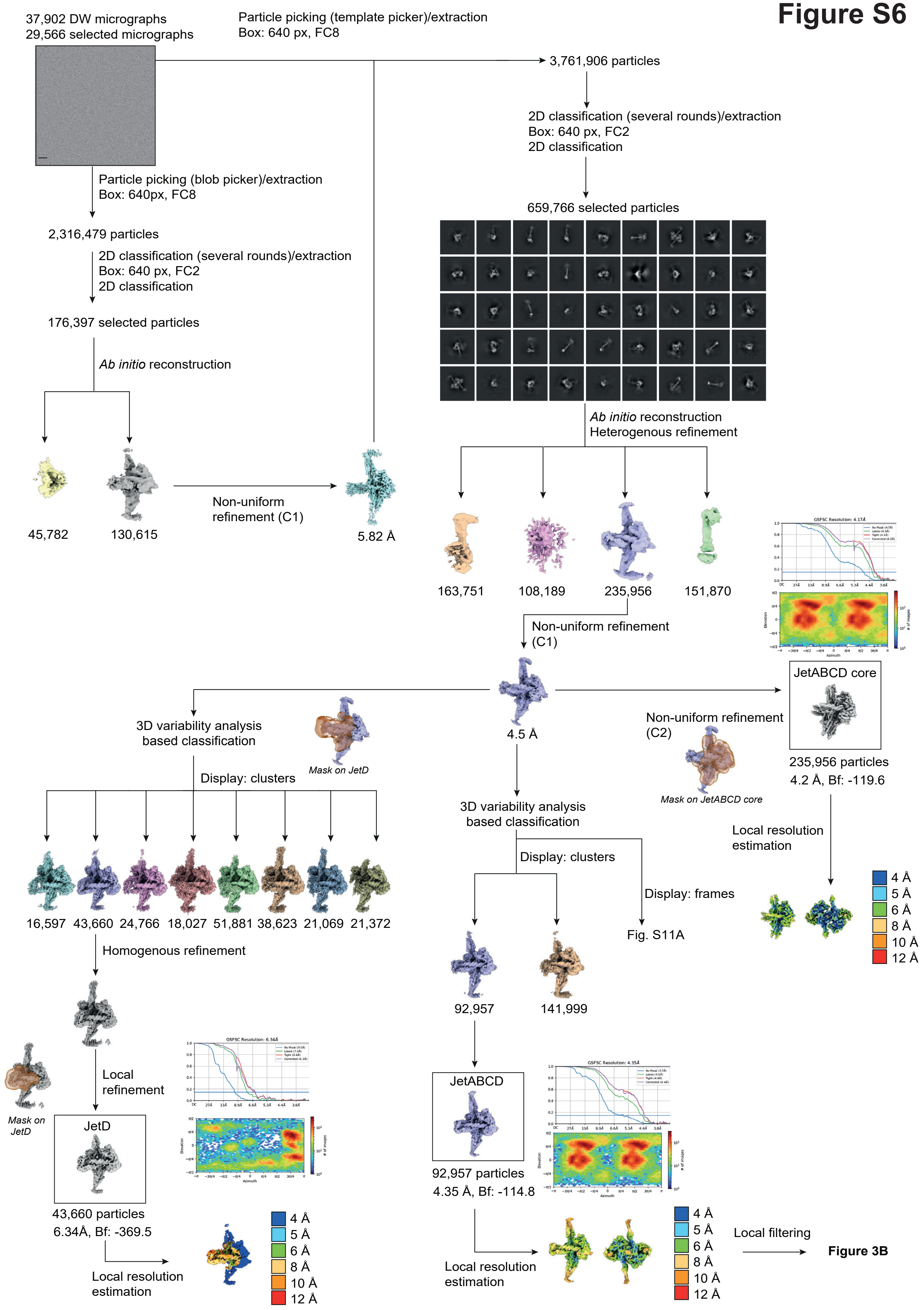

37,902 DW micrographs
29,566 selected micrographs
Particle picking (template picker)/extraction
Box: 640 px, FC8
3,761,906 particles
2D classification (several rounds)/extraction
Box: 640 px, FC2
2D classification
659,766 selected particles
Particle picking (blob picker)/extraction
Box: 640px, FC8
2,316,479 particles
2D classification (several rounds)/extraction
Box: 640 px, FC2
2D classification
176,397 selected particles
Ab initio reconstruction
45,782
130,615
Non-uniform
refinement (C1)
5.82 Å
Ab initio reconstruction
Heterogenous refinement
163,751
108,189
235,956
151,870
GSFSC Resolution: 4.17Å
Non-uniform refinement
(C1)
4.5 Å
JetABCD core
Non-uniform refinement
(C2)
Mask on JetABCD core
235,956 particles
4.2 Å, Bf: -119.6
3D variability analysis
based classification
Mask on JetD
Display: clusters
16,597
43,660
24,766
18,027
51,881
38,623
21,069
21,372
Homogenous refinement
3D variability analysis
based classification
Display: clusters
Display: frames
Fig. S11A
Local resolution
estimation
4 Å
5 Å
6 Å
8 Å
10 Å
12 Å
92,957
141,999
Local
refinement
Mask on
JetD
JetD
GSFSC Resolution: 6.34Å
43,660 particles
6.34Å, Bf: -369.5
Local resolution
estimation
4 Å
5 Å
6 Å
8 Å
10 Å
12 Å
JetABCD
GSFSC Resolution: 4.35Å
92,957 particles
4.35 Å, Bf: -114.8
Local resolution
estimation
4 Å
5 Å
6 Å
8 Å
10 Å
12 Å
Local filtering
Figure 3B

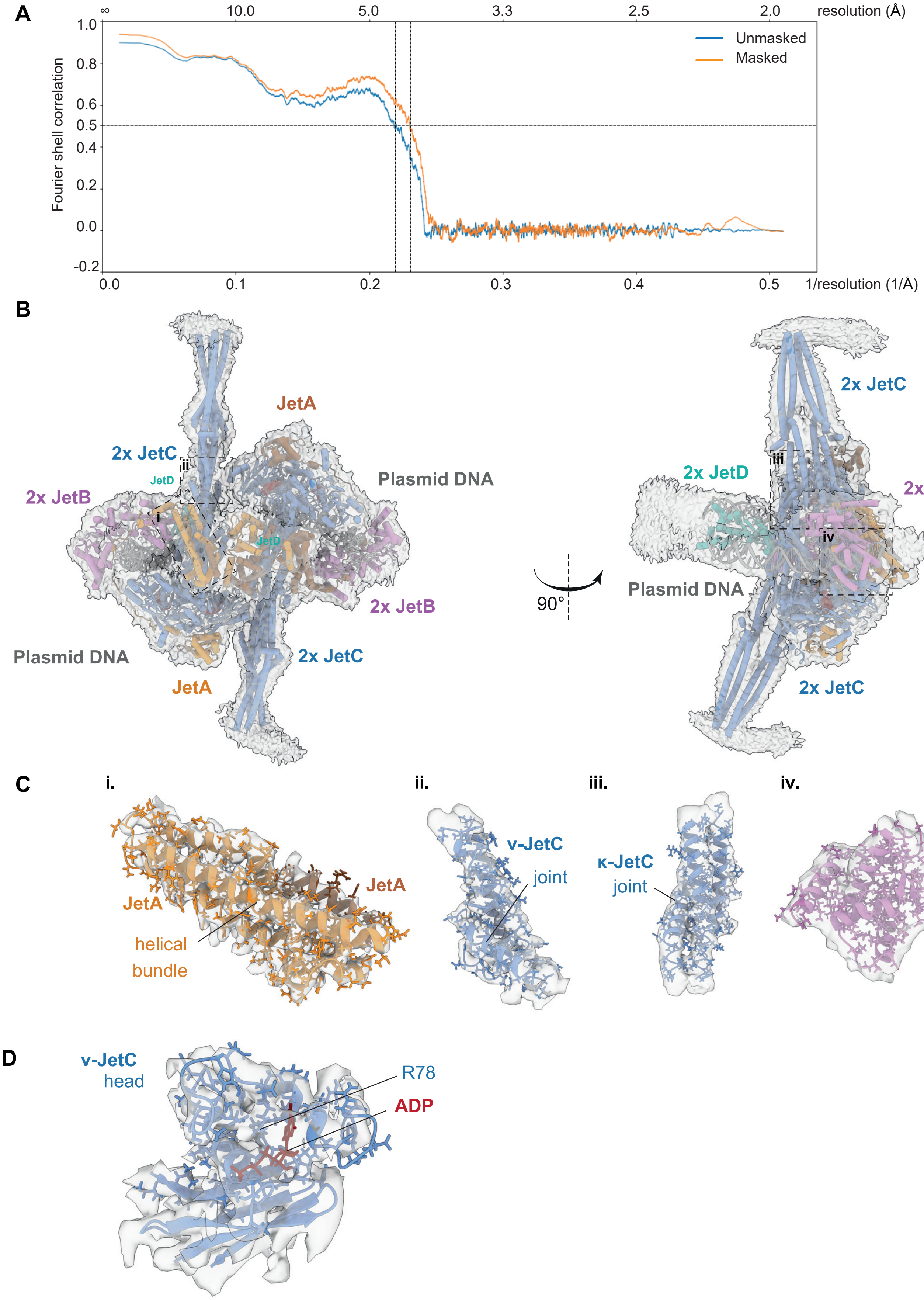

A
∞
10.0
5.0
3.3
2.5
2.0
resolution (Å)
1.0
0.8
0.6
0.5
0.4
0.2
0.0
-0.2
Fourier shell correlation
Unmasked
Masked
0.0
0.1
0.2
0.3
0.4
0.5
1/resolution (1/Å)
B
JetA
2x JetC
JetD
2x JetB
Plasmid DNA
2x JetB
Plasmid DNA
JetA
2x JetC
90°
2x JetC
2x JetD
2x JetB
2x JetA
Plasmid DNA
2x JetC
C
i.
ii.
iii.
iv.
JetA
JetA
helical
bundle
v-JetC
joint
κ-JetC
joint
JetB
cWHD
D
v-JetC
head
R78
ADP


### Figure S7

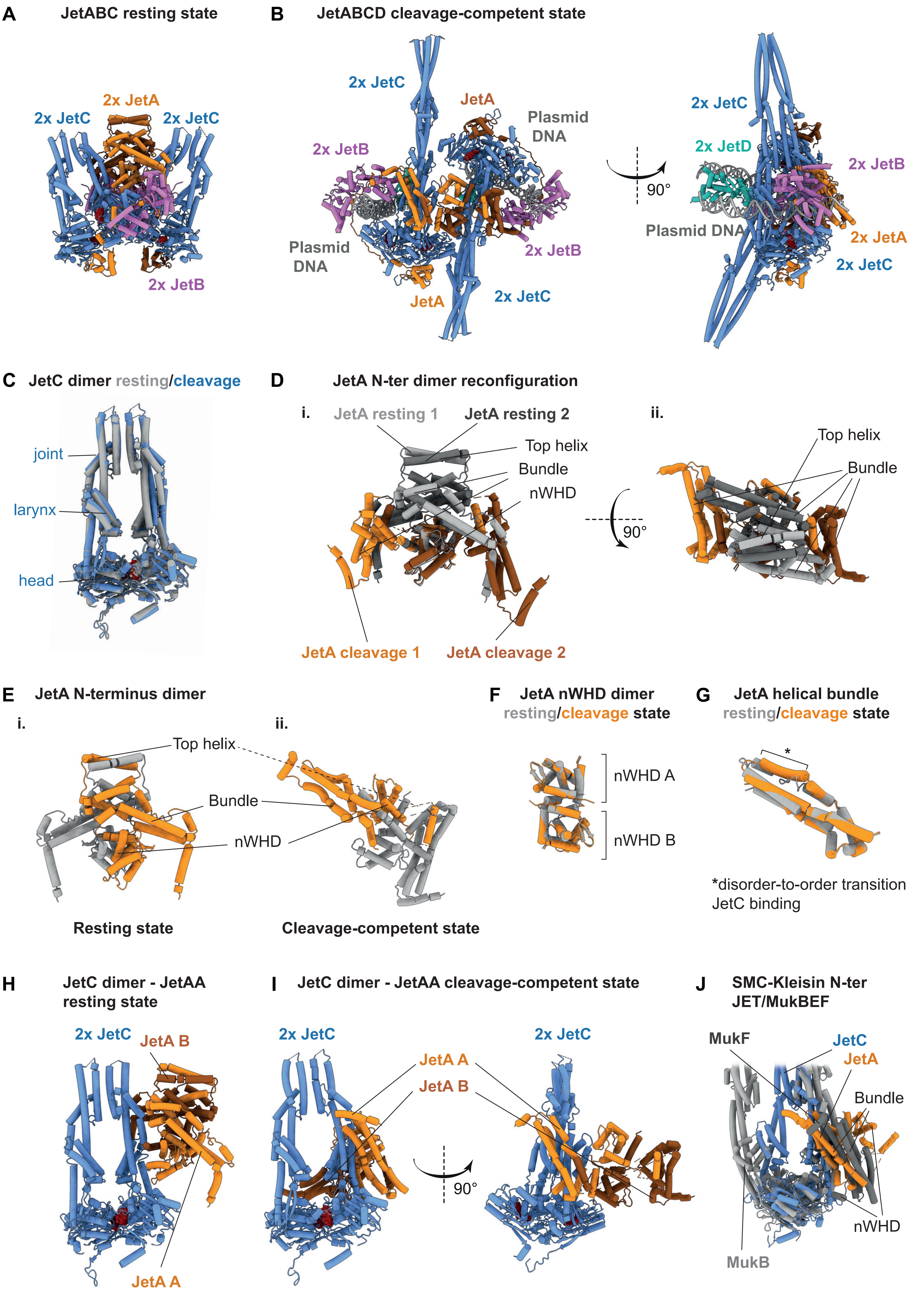
A
JetABC resting state
2x JetA
2x JetC
2x JetC
2x JetB
B
JetABCD cleavage-competent state
2x JetC
JetA
Plasmid DNA
2x JetB
Plasmid DNA
2x JetB
JetA
2x JetC
90°
2x JetC
2x JetD
2x JetB
Plasmid DNA
2x JetA
2x JetC
C
JetC dimer resting/cleavage
joint
larynx
head
D
JetA N-ter dimer reconfiguration
i.
JetA resting 1
JetA resting 2
Top helix
Bundle
nWHD
JetA cleavage 1
JetA cleavage 2
ii.
90°
Top helix
Bundle
E
JetA N-terminus dimer
i.
ii.
Top helix
Bundle
nWHD
Resting state
Cleavage-competent state
F
JetA nWHD dimer
resting/cleavage state
nWHD A
nWHD B
G
JetA helical bundle
resting/cleavage state
*
*disorder-to-order transition
JetC binding
H
JetC dimer - JetAA
resting state
2x JetC
JetA B
JetA A
I
JetC dimer - JetAA cleavage-competent state
2x JetC
JetA A
JetA B
90°
2x JetC
J
SMC-Kleisin N-ter
JET/MukBEF
MukF
JetC
JetA
Bundle
nWHD
MukB

### Figure S8

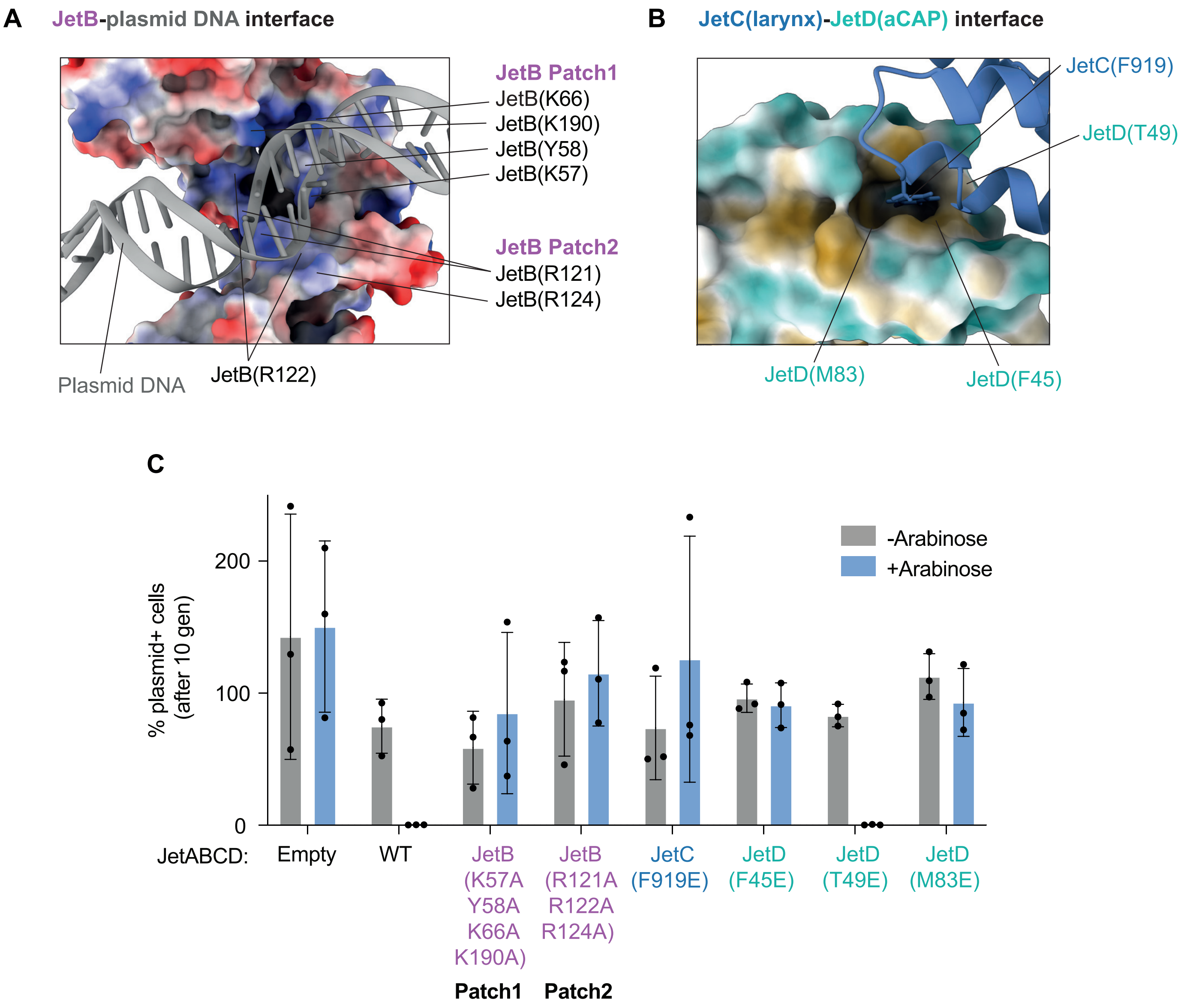
A
JetB-plasmid DNA interface
JetB Patch1
JetB(K66)
JetB(K190)
JetB(Y58)
JetB(K57)
JetB Patch2
JetB(R121)
JetB(R124)
Plasmid DNA
JetB(R122)
B
JetC(larynx)-JetD(aCAP) interface
JetC(F919)
JetD(T49)
JetD(M83)
JetD(F45)
C
% plasmid+ cells (after 10 gen)
200
100
0
-Arabinose
+Arabinose
JetABCD:
Empty
WT
JetB (K57A Y58A K66A K190A)
JetB (R121A R122A R124A)
JetC (F919E)
JetD (F45E)
JetD (T49E)
JetD (M83E)
Patch1
Patch2

### Figure S9

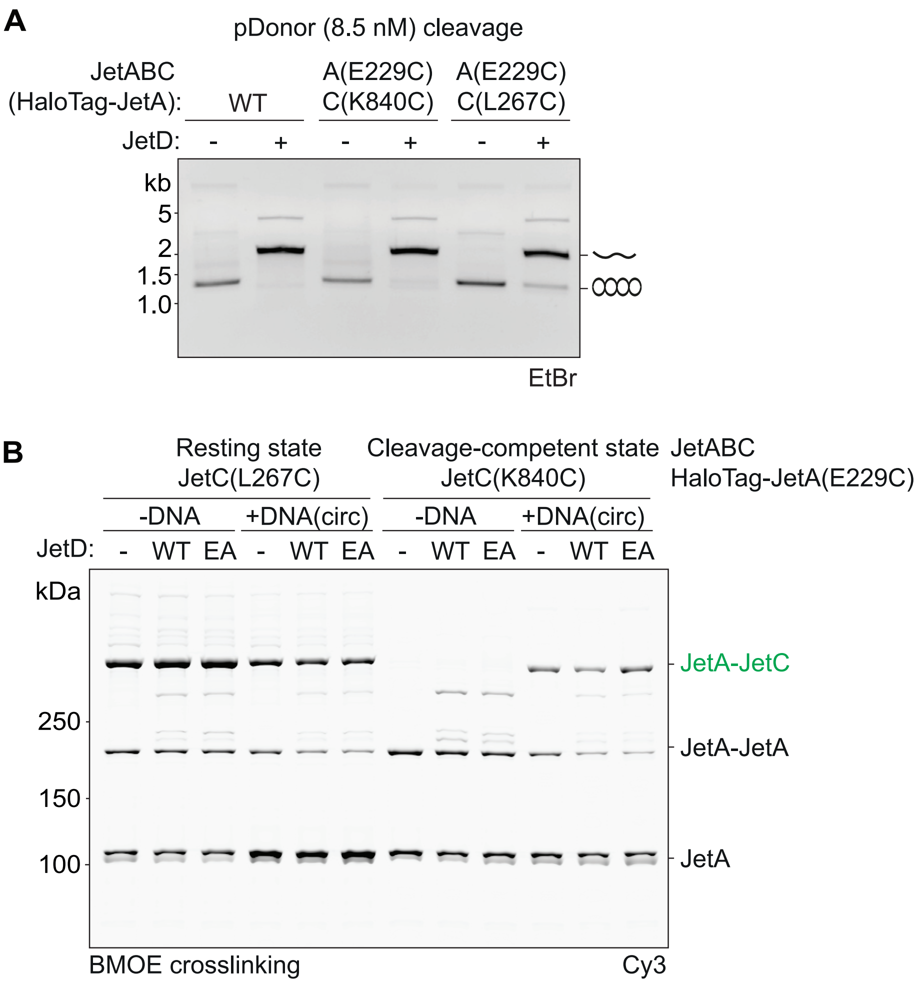
A
pDonor (8.5 nM) cleavage
JetABC
(HaloTag-JetA):
WT
A(E229C)
C(K840C)
A(E229C)
C(L267C)
JetD: - + - + - +
kb
5
2
1.5
1.0
EtBr
B
Resting state
JetC(L267C)
Cleavage-competent state
JetC(K840C)
JetABC
HaloTag-JetA(E229C)
-DNA
+DNA(circ)
-DNA
+DNA(circ)
JetD: - WT EA - WT EA - WT EA - WT EA
kDa
250
150
100
JetA-JetC
JetA-JetA
JetA
BMOE crosslinking
Cy3

### Figure S10

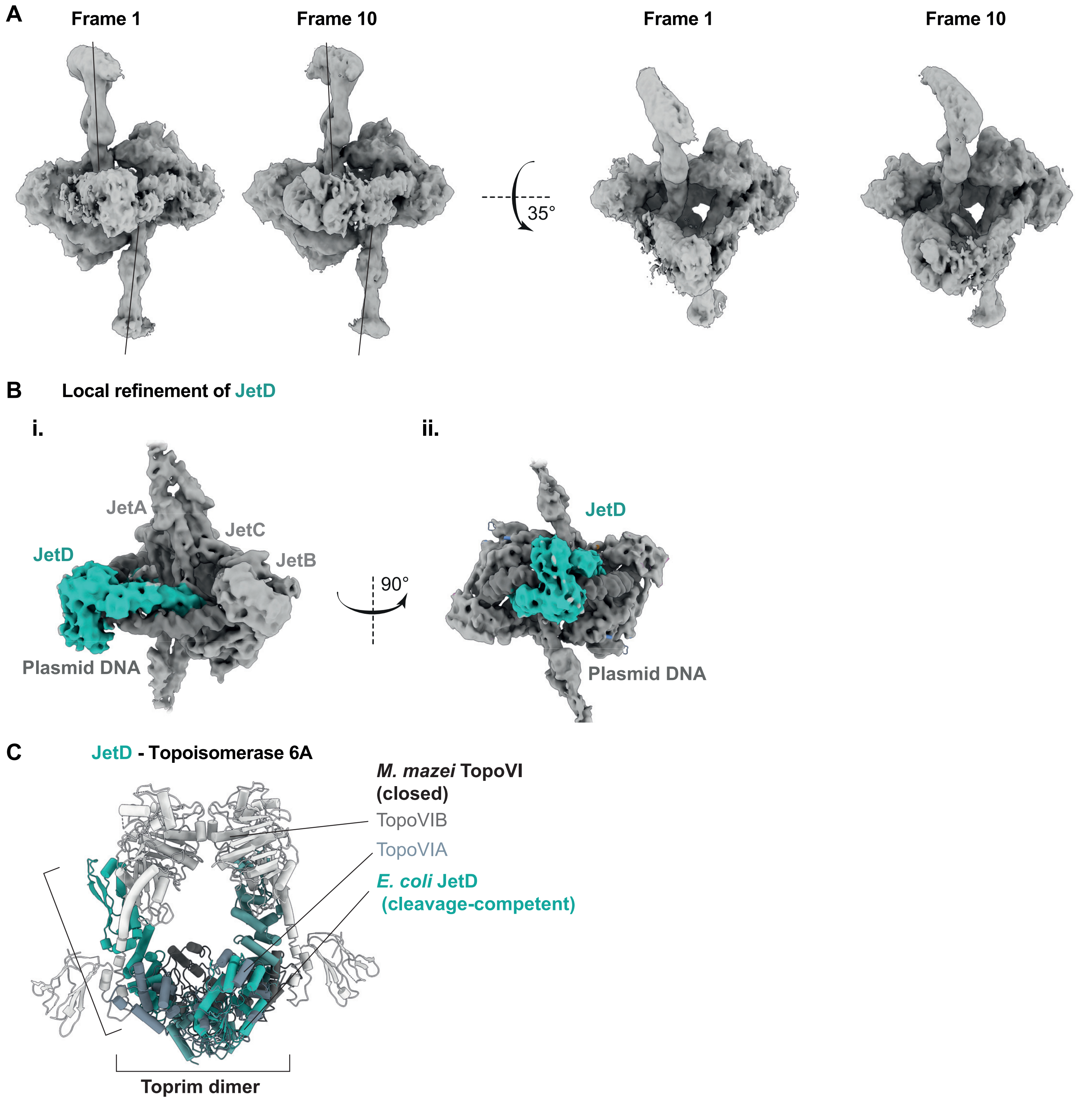
A
Frame 1
Frame 10
35°
Frame 1
Frame 10
B
Local refinement of JetD
i.
JetA
JetC
JetB
JetD
Plasmid DNA
90°
ii.
JetD
Plasmid DNA
C
JetD - Topoisomerase 6A
M. mazei TopoVI
(closed)
TopoVIB
TopoVIA
E. coli JetD
(cleavage-competent)
Toprim dimer

### Figure S11

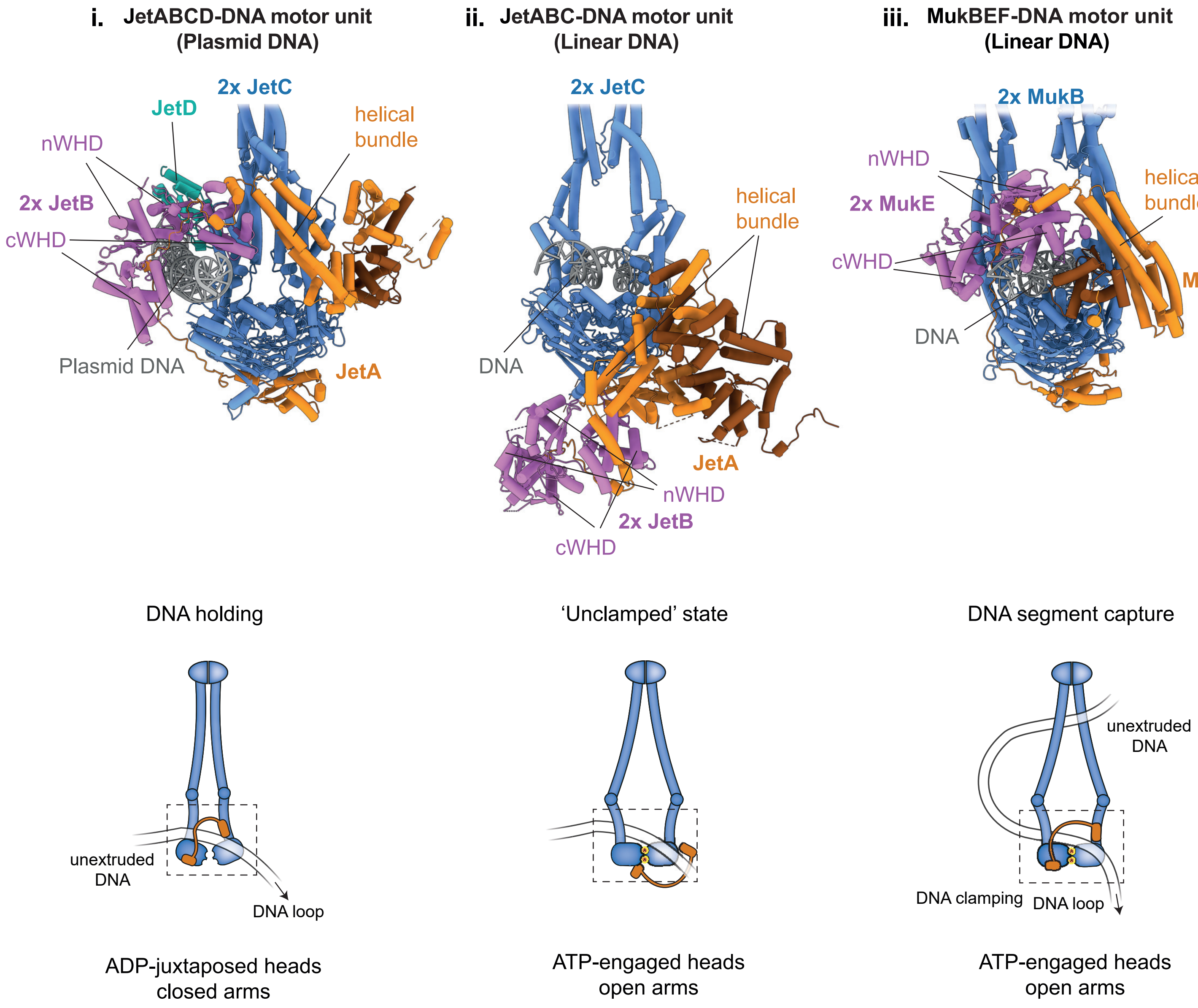
i. JetABCD-DNA motor unit (Plasmid DNA)
ii. JetABC-DNA motor unit (Linear DNA)
iii. MukBEF-DNA motor unit (Linear DNA)
2x JetC
JetD
helical bundle
nWHD
2x JetB
cWHD
Plasmid DNA
JetA
2x JetC
helical bundle
DNA
JetA
nWHD
2x JetB
cWHD
2x MukB
nWHD
2x MukE
helical bundle
cWHD
MukF
DNA
DNA holding
'Unclamped' state
DNA segment capture
unextruded DNA
DNA loop
unextruded DNA
DNA clamping
DNA loop
ADP-juxtaposed heads closed arms
ATP-engaged heads open arms
ATP-engaged heads open arms

### Figure S12
